# Protein Language Models Encode Evolutionary Grammar but Conflate Topological and Thermodynamic Phases

**DOI:** 10.64898/2026.04.07.717117

**Authors:** Yiquan Wang, Minnuo Cai, Yahui Ma, Xu Wang, Kai Wei

## Abstract

How a one-dimensional sequence encodes a dynamic thermodynamic ensemble rather than a static geometry remains a fundamental biophysical challenge. While protein language models achieve high structural accuracy from single sequences, it remains unclear whether they internalize physical folding principles or merely capture evolutionary statistical correlations. We investigate this representational capacity by analyzing the latent manifold of ESM-2 across 11068 proteins, focusing on fundamental exceptions to the Anfinsen assumption: intrinsically disordered, fold-switching, and knotted proteins. We demonstrate that ESM-2 integrates out microscopic backbone geometry during feature extraction. It instead forms a macroscopic sequence grammar manifold that separates random sequences while ordering biological proteins along physicochemical composition gradients. Consequently, the model exhibits topological aliasing. It conflates these physically distinct categories because they frequently share overlapping sequence statistics despite divergent three-dimensional topologies. A region-replacement experiment establishes this aliasing as an intrinsic characteristic rather than an artifact of sequence dilution, and a structure-aware control model (SaProt) demonstrates that explicit structural tokens partially alleviate topological blindness for static anomalies while failing to resolve multi-state thermodynamic phases. Furthermore, local curvature analysis reveals class-invariant geometric turbulence that homogenizes microscopic details while preserving macroscopic distinguishability. ESM-2 therefore functions as an evolutionary grammar compressor rather than a microscopic threedimensional geometry encoder, imposing defined boundaries on tasks requiring high-precision geometric awareness and necessitating integration with explicit physical constraints.

## 1 Introduction

Proteins are dynamic thermodynamic ensembles that explore conformational landscapes shaped by competing physical interactions rather than static geometric objects. Understanding how macroscopic folding behavior emerges from microscopic sequence information remains a central challenge in biophysics. Recent advances in structure prediction achieve high accuracy without explicit thermodynamic supervision. The capacity of these models to predict structure prompts an examination of whether their latent spaces encode physical geometry or organize sequences by evolutionary vocabulary. We analyze ESM-2 [1,2] to investigate this distinction. ESM2 maps single sequences directly to latent representations and provides a purely sequence driven framework isolated from the explicit spatial and co-evolutionary contact signals introduced by multiple sequence alignments in systems like AlphaFold3 [3, 4]. Exploring the continuous latent spaces of deep generative models has indeed proven highly effective for mapping evolutionary relationships and optimizing sequence fitness landscapes [5, 6]. The success of structure prediction tools relies heavily on downstream equivariant modules mapping latent representations to known spatial coordinates. This architectural division obscures the intrinsic geometric content of the latent space itself. Distinguishing the encoded information requires evaluating the models against proteins that explicitly violate the classical Anfinsen assumption of one sequence mapping to one stable structure [7].

For standard proteins evolutionary statistics and physical folding rules exhibit a strong correlation. Intrinsically disordered proteins fold-switching proteins and knotted proteins (Figure 1) provide the necessary test cases to probe the physical awareness of language models because they decouple this statistical correlation from physical reality. These three categories represent fundamental structural exceptions to the Anfinsen paradigm. IDPs possess a single sequence but lack fixed geometric structure to exhibit continuous disorder and mediate regulatory networks through phase separation [8–12]. Although recent language models and generative approaches can predict disorder and model dynamic ensembles [13, 14], IDP functions are fundamentally mediated by conserved evolutionary features rather than static geometry [15]. Indeed, protein language models and deep generative frameworks successfully capture this evolutionary grammar and specific compositional motifs of IDP phase separation despite their extreme sensitivity to dynamic chemical environments [16–19]. Fold-switching proteins adopt multiple discrete stable folds from a single sequence in response to cellular stimuli and thermodynamic balances [20], and challenge the foundation of static structure prediction [21–23]. These proteins exploit fundamental ambiguities in the folding code [24] and possess distinct co-evolutionary signatures for both conformations that are masked in standard sequence alignments [25]. Knotted proteins feature backbones forming non-trivial topological knots that alter folding kinetics [26–28] while remaining invisible to standard sequence composition statistics despite their strict evolutionary conservation across divergent families [29, 30]. Extending topological data analysis, which has recently provided global structural classifications and representation learning for the physical biomolecular universe [31, 32], to the neural latent space determines whether protein language models encode physical spatial constraints or capture evolutionary amino acid combinations.

**Figure 1.**
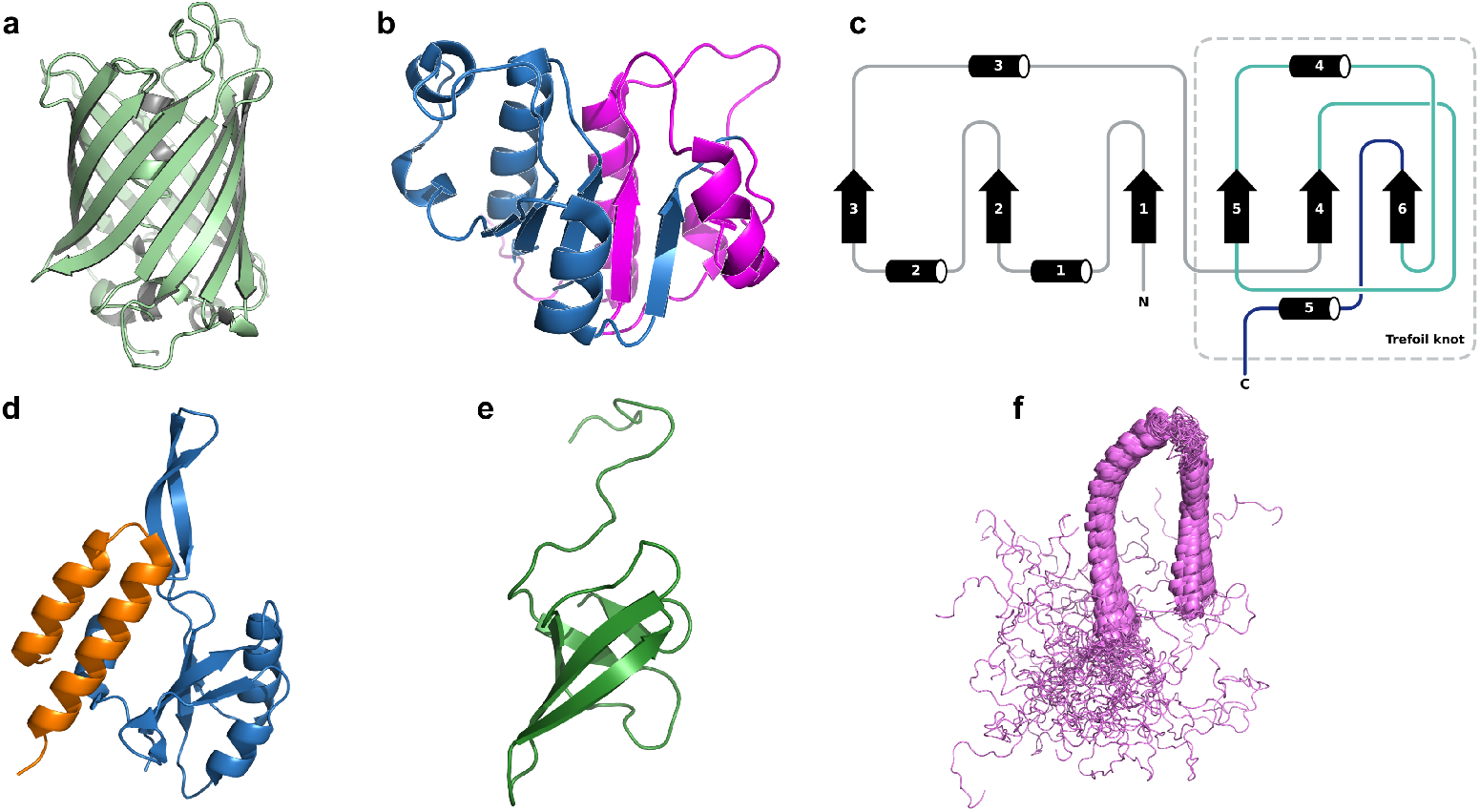
Physical and topological exceptions to the classical Anfinsen assumption. Representative three-dimensional structures illustrating the fundamental divide between rigid geometry and thermodynamic/topological anomalies. **(a)** A standard rigid protein: green fluorescent protein (GFP; PDB: 1GFL), displaying a single, stable *β*-barrel fold characteristic of the Anfinsen paradigm. **(b)** A knotted protein: YibK methyltransferase (PDB: 1MXI), colored by topological role—sky blue for the rigid core (residues 1–80) and magenta for the deep trefoil (3_1_) knot region (residues 81–160), in which the C-terminal chain threads through a loop formed by the preceding segment. **(c)** Topology diagram of YibK, illustrating the secondary-structure connectivity and the trefoil-knot motif (cyan box). Arrows denote *β*-strands and cylinders denote *α*-helices; N and C mark the chain termini. **(d)** Fold-switching protein RfaH in the autoinhibited *α*-helical state (PDB: 5OND, chain A). The NTD (residues 1–99, blue) and the KOW/CTD domain (residues 115–162, orange) are shown; the CTD adopts a two-helix hairpin. **(e)** The same KOW domain of RfaH in the activated *β*-barrel state (PDB: 2LCL, green), demonstrating that an identical sequence can adopt two entirely distinct stable folds. **(f)** An intrinsically disordered protein: *α*-synuclein (PDB: 2KKW), shown as an overlay of 34 NMR conformers (violet) obtained in the presence of SLAS micelles. The N-terminal region retains partial *α*-helical character while the C-terminal tail samples a broad conformational ensemble, highlighting the absence of a unique ground-state geometry. All structures are rendered in cartoon representation; solvent and heteroatoms have been removed for clarity.

The information encoded by protein language models occupies two possible physical scales. Microscopic encoding requires the model to represent precise local geometric details including exact dihedral angles and hydrogen bond networks. Macroscopic encoding implies the model abstracts global evolutionary properties and physicochemical composition. We leverage the Hasimoto integrability error *E*[*n*] as a rigorous differential geometric quantity to distinguish these scenarios. The Hasimoto map [33] translates continuous three-dimensional space curves into mathematical solutions of the discrete nonlinear Schrödinger equation. Recent physical frameworks establish that the residual of this mapping, the quantity *E*[*n*], functions as a precise kinematic identity and geometric order parameter for the microscopic threedimensional twisting and spatial folding of the protein backbone [34, 35]. Furthermore, this discrete geometric invariant rigorously quantifies sub-residue structural phase transitions and the mathematical characterization of biological order [36, 37]. Correlating latent space embeddings with *E*[*n*] determines whether the model encodes the exact geometric reality of the atomic coordinates or operates on sequence-level statistical approximations.

We analyze the latent manifold of ESM-2 across 11,068 protein sequences using a combined framework of differential geometry, topological data analysis, and gauge field theory. The dataset spans seven functional categories to ensure comprehensive structural diversity. These categories include ASTRAL95 structural representatives, integrable helices, random sequences, IDPs, fold-switching proteins, and knotted proteins. We establish a scale separation where microscopic three-dimensional backbone geometry is decoupled from the macroscopic latent manifold. The latent space functions as a sequence grammar manifold that maps evolutionary statistics onto physicochemical composition gradients to isolate random sequences from biological ones, a fundamental division functionally mirrored in the topology of physical energy landscapes [38]. This statistical reliance results in topological aliasing. The model conflates IDPs, knotted proteins, and fold-switching proteins because these physically disparate categories frequently share overlapping sequence statistics. A region-replacement experiment provides compelling evidence against the hypothesis that this aliasing arises from mathematical dilution of distinctive domains and establishes it as an intrinsic limitation of the representation. The latent space of ESM-2 encodes a macroscopic evolutionary sequence grammar but fails to resolve the microscopic topological and thermodynamic phases of physical protein folding.

## 2 Results

### 2.1 Microscopic Geometry Is Absent from the Latent Manifold

We first evaluated the encoding of microscopic backbone geometry by analyzing 9195 protein sequences from the anchor astral95 and integrable categories with computable Hasimoto integrability error *E*[*n*]. We computed the residue-level Spearman correlation between embedding distances and *E*[*n*] along the backbone for each protein.

Figure 2a demonstrates that residue-level correlations are negligible with an overall Spearman *ρ* = 0.1053 (*p* = 4.18 × 10^−24^) a median |*ρ*| = 0.1147 and a linear fit yielding *R* ^2^ = 0.0152. The scatter pattern exhibits isotropic noise largely devoid of visually coherent signal. The 9195 sequences with *E*[*n*] show no visible gradient across the horseshoe structure when projected onto the PCA manifold (Figure 2b) demonstrating *ρ*(PC1, *E*[*n*]) = −0.150 (*p* = 2.3 × 10^−47^) and *ρ*(PC2, *E*[*n*]) = 0.140 (*p* = 3.4 × 10^−41^). This confirms that macroscopic manifold organization operates independently of microscopic geometry.

**Figure 2.**
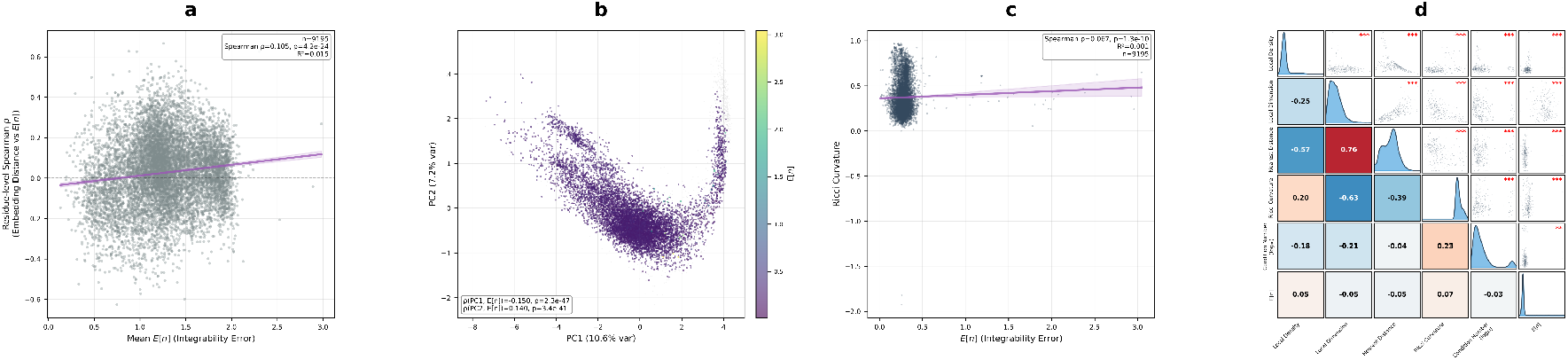
Scale separation in the ESM-2 latent space. We analyzed 9,195 protein sequences with valid Hasimoto integrability error *E*[*n*] derived from the anchor, astral95, and integrable datasets. **(a)** The residue level Spearman *ρ* comparing embedding distance against *E*[*n*] is plotted alongside the mean *E*[*n*] for each protein. The purple line represents the linear fit and the shaded region denotes the 95 percent confidence interval. **(b)** The PCA projection incorporates all 11,068 sequences in the dataset. Sequences possessing a computed *E*[*n*] are colored according to their *E*[*n*] value using a viridis colormap. The remaining 1,873 sequences lacking an *E*[*n*] definition are displayed in light gray. The first principal component explains 10.60 percent of the total variance and the second principal component explains 7.20 percent. **(c)** The scatter plot visualizes *E*[*n*] relative to Ollivier-Ricci curvature. The purple line delineates the linear fit accompanied by a 95 percent confidence interval. **(d)** The Spearman correlation matrix evaluates six geometric variables including Local Density, Local Dimension, Nearest Distance, Ricci Curvature, log_10_ Condition Number, and *E*[*n*]. The matrix diagonal displays kernel density estimates. The lower triangle presents a correlation coefficient heatmap using an RdBu colormap. The upper triangle contains scatter plots enriched with significance annotations where ^∗∗∗^ indicates *p <* 0.001, ^∗∗^ indicates *p <* 0.01, and ^∗^ indicates *p <* 0.05. Local geometric features were computed from 20 nearest neighbors within the 50 dimensional PCA space. All statistical tests utilize two tailed Spearman rank correlations.

This structural decoupling extends to intrinsic manifold geometry. The correlation between *E*[*n*] and Ollivier-Ricci curvature [39] yields a Spearman *ρ* = 0.067 (*p* = 1.3 × 10^−10^ Figure 2c). A comprehensive correlation matrix (Figure 2d) confirms that *E*[*n*] shows |*ρ*| *<* 0.07 with all five latent geometric features including Local Density Local Dimension Nearest Distance Ricci Curvature and Condition Number. These features remain strongly inter-correlated among themselves with *ρ*(Local Dimension, Nearest Distance) = 0.760. Category-decoupled analysis (Figure S2) verifies that this decoupling holds within each protein class and rules out the Simpson paradox.

The observed scale separation indicates that the Transformer architecture of ESM-2 performs a coarse-graining operation that averages out microscopic three-dimensional geometry. The mean-pooling operation integrates out highfrequency geometric fluctuations to yield a macroscopic representation organized by evolutionary sequence statistics rather than local backbone geometry. The precise spatial arrangements required for catalytic active sites or specific drug-binding pockets are consequently lost in the latent space. This characteristic limits the utility of this global representation for atomic-resolution molecular design without external structural refinement.

### 2.2 The Latent Manifold Organizes by Sequence Grammar

The absence of microscopic geometric encoding directs the analytical focus toward the macroscopic topology of the latent space. The PCA projection of all 11068 sequences reveals this structural architecture as a characteristic horseshoe-shaped manifold (Figure 3a) where PC1 explains 10.60% and PC2 explains 7.20% of the variance.

**Figure 3.**
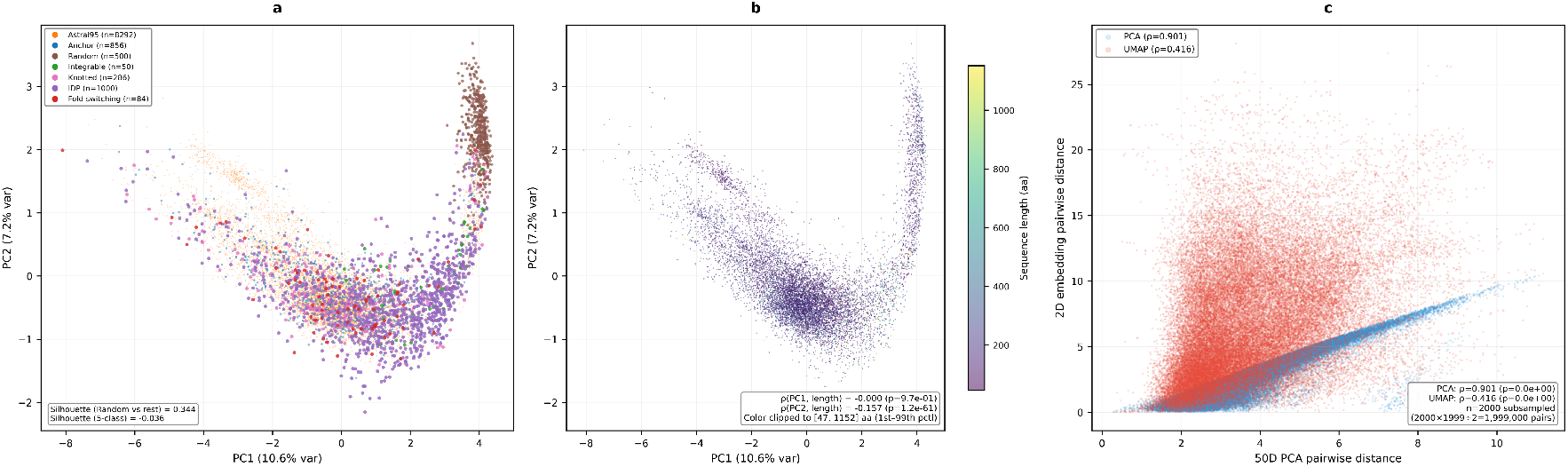
The sequence grammar manifold. We projected 11,068 protein sequences from the ESM-2 latent space using principal component analysis. **(a)** The two dimensional PCA projection is colored according to seven distinct protein categories. The first and second principal components explain 10.60 percent and 7.20 percent of the variance respectively. The manifold forms a defined horseshoe distribution. Random sequences shown in brown are robustly isolated at the right tip of the manifold. Non random biological categories merge into a highly mixed main body. All legend markers maintain a uniform size for clarity. **(b)** The PCA projection is colored by sequence length using a viridis colormap clipped between the 1st and 99th percentiles spanning 47 to 1152 amino acids to suppress extreme outliers. The orthogonal direction of the color gradient confirms that the first principal component does not trivially correlate with sequence length. **(c)** We compared pairwise distance preservation between the two dimensional PCA projection and the two dimensional UMAP projection relative to the 50 dimensional PCA space. The scatter plots display 50,000 distance pairs randomly sampled from the dataset. The PCA algorithm demonstrates significantly stronger preservation of global pairwise distances when compared directly to UMAP.

The most prominent structural feature is the definitive geometric isolation of random sequences. The 500 random sequences cluster tightly at the right tip of the horseshoe and achieve a Silhouette Score [40] of 0.344 using Euclidean distance in the 50D PCA space. This pattern indicates that ESM-2 internalizes the statistical regularities of evolved sequences and reliably differentiates them from random amino acid combinations. The overall 5-class Silhouette Score is - 0.036 which indicates that non-random categories are highly intermixed within the manifold body. The model successfully distinguishes unphysical polymers but organizes valid biological proteins by their shared evolutionary vocabulary rather than partitioning them into rigid structural bins.

The principal axis PC1 does not function as a trivial length counter. Sequence lengths in our dataset range from 10 to 7073 amino acids with a mean of 218.9 *±* 240.1 aa and a median of 159 aa. Coloring the projection by sequence length (Figure 3b) reveals *ρ*(PC1, length) = −0.0003 (*p* = 9.74 × 10^−1^) although PC2 shows a weak negative correlation *ρ* = −0.157 (*p* = 1.21 × 10^−61^). We validated the choice of PCA as the primary coordinate system by comparing distance preservation between PCA-2D and UMAP-2D relative to the 50D PCA space (Figure 3c). Using 1999000 pairwise Euclidean distances generated from a random subsample of 2000 sequences PCA-2D achieves a Spearman *ρ* = 0.901 (*p* ≈ 0) while UMAP-2D achieves *ρ* = 0.416 (*p* ≈ 0). PCA preserves superior global distance structure and serves as the optimal basis for geometric analysis while UMAP provides advantageous local structure emphasis for density estimation (Figure S3 and Table S1).

### 2.3 Latent Principal Axes Capture Physicochemical Properties

The geometric isolation of random sequences verifies that the manifold encodes fundamental evolutionary constraints. Mapping specific physicochemical properties and SCOP structural classes onto the PCA projection translates these mathematical axes into concrete biological determinants (Figure 4). PC2 emerges as a biochemical composition axis that correlates moderately with GRAVY hydropathy [41] (*ρ* = − 0.214 *p* = 7.88 × 10^−115^ Figure 4a) weakly with isoelectric point pI (*ρ* = 0.153 *p* = 2.11 × 10^−58^ Figure 4b) and most strongly with aromaticity (*ρ* = 0.364 Figure 4d). The opposing signs of the GRAVY and pI correlations with PC2 establish a complementary hydropathy gradient where the upper arm of the horseshoe is hydrophilic and the lower arm is hydrophobic.

**Figure 4.**
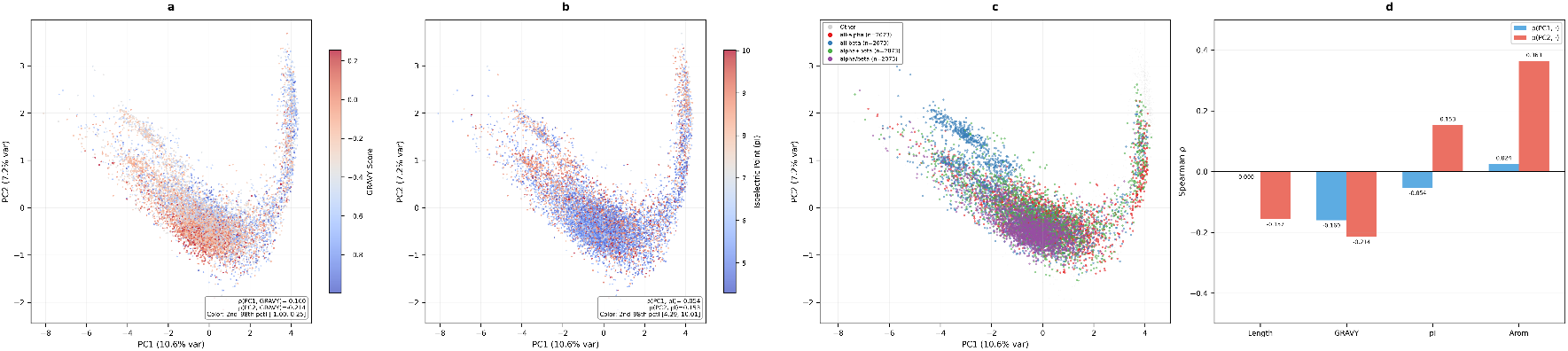
Decoding the latent axes. We mapped physicochemical properties and structural classes onto the PCA projection of 11,068 protein sequences from the ESM-2 latent space. **(a)** The PCA projection is colored by the GRAVY hydropathy index for 11,032 valid sequences. The visualization employs a coolwarm colormap clipped between the 2nd and 98th percentiles spanning −1.00 to 0.26. **(b)** The projection is colored by the isoelectric point pI using a coolwarm colormap clipped between the 2nd and 98th percentiles spanning 4.29 to 10.01. **(c)** The projection is partitioned by SCOP structural class using the ASTRAL95 subset. The all-alpha class comprises 2,073 sequences marked in red. The all-beta class comprises 2,073 sequences marked in blue. The alpha+beta class comprises 2,073 sequences marked in green. The alpha/beta class comprises 2,073 sequences marked in purple. The remaining 2,776 sequences appear in light gray. Legend markers maintain uniform scaling. **(d)** The bar chart summarizes Spearman rank correlations. We evaluated four properties including Sequence Length, GRAVY, pI, and Aromaticity against the first principal component shown in blue and the second principal component shown in red. All statistical comparisons rely on two tailed Spearman rank correlations.

PC1 resists reduction to any single physicochemical property and yields |*ρ*(PC1, ·)| *<* 0.17 for all four properties tested (Figure 4d). PC1 shows only weak associations with GRAVY (*ρ* = −0.160 *p* = 4.59 × 10^−64^) and pI (*ρ* = −0.054 *p* = 1.88 × 10^−8^) which suggests it encodes a complex sequence grammar integrating multiple evolutionary signals rather than simple compositional metrics.

SCOP structural classes [42] show ordered arrangement −0.031. along the horseshoe (Figure 4c) where all-alpha all-beta alpha+beta and alpha/beta classes occupy distinct but overlapping regions. This structural ordering indicates that the manifold reflects secondary structure preferences as a statistical property of sequences rather than encoding detailed three-dimensional coordinates.

### 2.4 Topological Aliasing Represents an Intrinsic Limitation

A representation governed strictly by one-dimensional sequence statistics faces an intrinsic challenge when evaluating proteins that violate the classical one sequence to one structure paradigm. Testing the model against IDPs knotted proteins and fold-switching proteins directly probes this representational boundary. These categories share standard evolutionary sequence statistics while diverging in three-dimensional topology and thermodynamic phase behavior. They effectively decouple the evolutionary sequence grammar from the physical folding conformation.

We computed per-sample Silhouette Scores in the PCA 50D space using the 7 class label system to quantify category separability (Figure 5). The results reveal a pronounced asymmetry where random sequences achieve a strongly positive score of *µ* = +0.566 (*n* = 500) and confirm reliable grammar discrimination. All three categories with non-trivial topology or thermodynamic properties exhibit negative scores including Knotted (*µ* = −0.151 *±* 0.133 *n* = 286) Fold-switching (*µ* = −0.108 *±* 0.129 *n* = 84) and IDP (*µ* = −0.057 *±* 0.119 *n* = 1000). These negative values indicate that intra-cluster distances exceed inter-cluster distances and establish a clear pattern of representational aliasing (Figure 5a and c). The overall 7 class Silhouette Score remains at −0.031.

**Figure 5.**
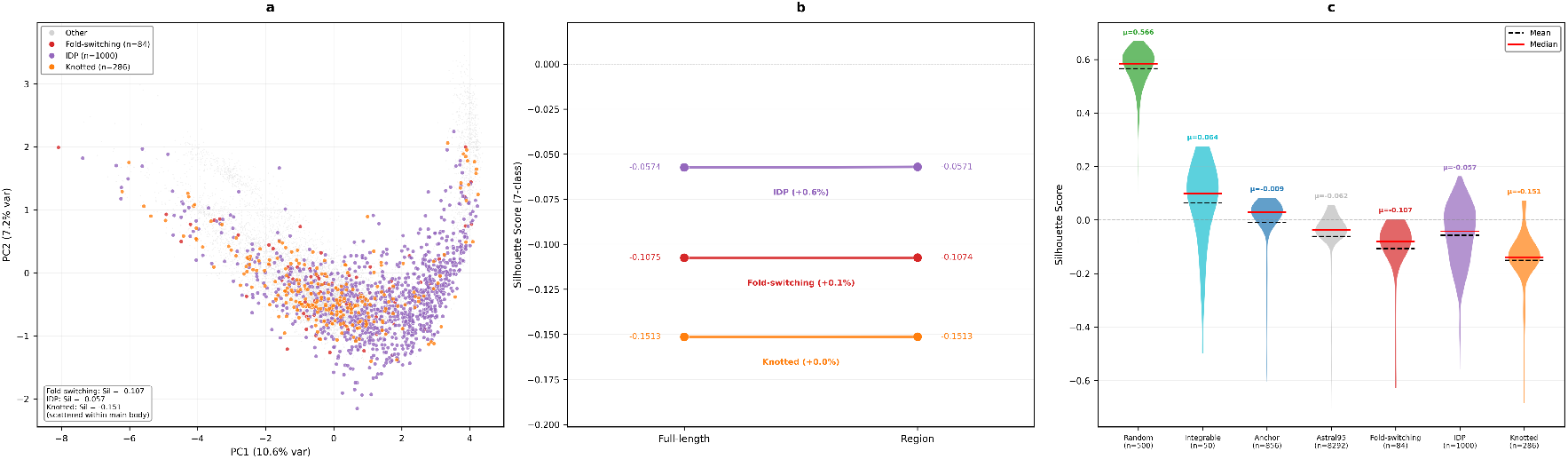
Topological aliasing and region replacement analysis. We quantified category separability for 11,068 protein sequences within the ESM-2 latent space utilizing a seven class Silhouette Score framework covering Random, Integrable, Anchor, Astral95, Fold-switching, IDP, and Knotted proteins. **(a)** The two dimensional PCA projection highlights the spatial distribution of three conformationally extreme protein categories. Fold-switching proteins are marked in red, intrinsically disordered proteins are marked in purple, and knotted proteins are marked in orange. These samples remain entirely dispersed within the light gray main manifold body composed of Astral95 and Anchor sequences. All three extreme categories record negative seven class Silhouette Scores. The Fold-switching score is −0.108 *±* 0.129, the IDP score is −0.057 *±* 0.119, and the Knotted score is −0.151 *±*0.133 based on Euclidean distances in the region replaced 50 dimensional PCA space. **(b)** The slope chart tracks Silhouette Score deviations between full length and region replaced embeddings. We isolated the three subcategories possessing explicit region annotations including fold-switching interface units, disordered regions, and knotted regions. Each independent line connects the full length score on the left axis to the region replaced score on the right axis. The Fold-switching mean score transitions from −0.1075 to −0.1074 reflecting a +0.1 percent shift. The IDP score transitions from −0.0574 to −0.0571 reflecting a +0.6 percent shift. The Knotted score remains stable at −0.1513 reflecting exactly a +0.0 percent shift. These near horizontal trajectories confirm that local distinctive regions do not drive category separability. **(c)** The violin plot illustrates per sample Silhouette Scores across all seven functional categories derived from the region replaced 50 dimensional PCA space. The categories are ordered by mathematical mean from highest to lowest. Random sequences lead with *µ* = +0.566 followed by Integrable sequences with *µ* = +0.064. Anchor sequences drop to *µ* = −0.009 and Astral95 sequences reach *µ* = −0.062. Fold-switching sequences record *µ* = −0.108, IDP sequences record *µ* = −0.057, and Knotted sequences conclude at *µ* = −0.151. Black dashed lines mark the statistical means and red solid lines indicate the medians. The gray dashed vertical line denotes the zero score boundary indicating complete clustering overlap.

This representational aliasing could potentially arise from a mathematical dilution effect where distinctive local regions are averaged out by flanking domains during the mean-pooling operation. We performed a targeted region-replacement experiment to test this sequence dilution hypothesis. The distinctive regions of each extreme protein were replaced with the corresponding regions from a matched ASTRAL95 protein and Silhouette Scores were recomputed (Figure 5b). The results reveal near zero changes in separability with Fold-switching scores changing by +0.1 percent IDP by +0.6 percent and Knotted by +0.0 percent. These minor variations produce near horizontal lines in the slope chart and indicate that the topological conflation does not arise from sequence dilution. This phenomenon constitutes an intrinsic representational characteristic of the sequence based framework where the model integrates physical geometric features into broad statistical categories (Figure S4). Proteins violating the classical structure paradigm map to highly overlapping regions with static rigid folds when their evolutionary sequence compositions are similar. This representational overlap explains the fundamental difficulty of utilizing sequence only embeddings to predict dynamic physical properties such as allosteric regulation disorder driven phase separation and kinetic trapping in knotted topologies.

### 2.5 Latent Density Inverts Physical Conformational Entropy

The geometric conflation of these topological anomalies necessitates an examination of their collective spatial distribution within the shared latent space. Manifold density analysis reveals a structural pattern of density inversion where functionally extreme proteins occupy systematically denser regions of the latent space than the broad structural reference set rather than dispersing as representational noise (Figure 6a). The three-dimensional density landscape exhibits a smooth core-to-periphery terrain gradient with a log_10_ density range from −3.71 to −1.63.

**Figure 6.**
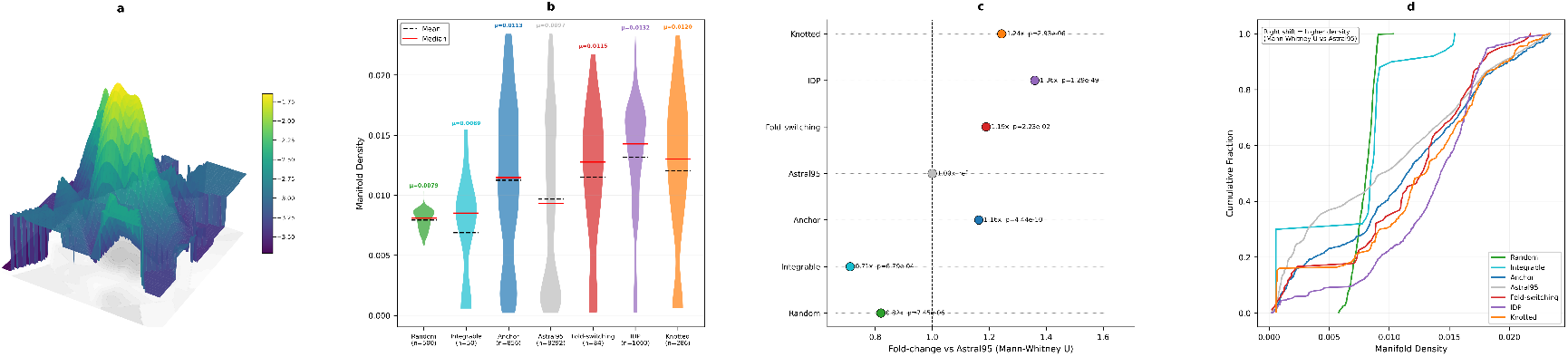
Density inversion in the ESM-2 latent space. Manifold density was estimated for 11,068 protein sequences across seven functional categories using kernel density estimation (KDE) on UMAP-2D coordinates. **(a)** Three-dimensional density landscape over the UMAP plane, represented as a surface colored by log_10_ density. **(b)** Violin plot of manifold density distributions. Black dashed lines indicate means, and red solid lines indicate medians; *µ* values are annotated above each violin. **(c)** Cleveland dot plot showing the density fold-change of each category relative to the broad structural reference set (Astral95). Exact *p*-values from Mann-Whitney *U* tests are annotated. **(d)** Cumulative distribution function (CDF) curves of manifold density. Rightward displacement indicates higher density relative to the Astral95 baseline.

Kernel density estimation on UMAP-2D coordinates using the Scott rule bandwidth demonstrates a strict density rank order (Figure 6b and Methods) with IDP at *µ* = 0.01316 Knotted at *µ* = 0.01203 Fold-switching at *µ* = 0.01150 Anchor at *µ* = 0.01125 Astral95 at *µ* = 0.00968 Random at *µ* = 0.00793 and Integrable at *µ* = 0.00689. IDP density reaches 1.36× that of the Astral95 baseline (*p* = 1.29*×*10^−49^ Mann-Whitney *U* ) Knotted reaches 1.24× (*p* = 2.93 × 10^−6^) and Fold-switching reaches 1.19× (*p* = 2.23 × 10^−2^ Figure 6c). Random sequences are conversely 0.82× less dense (*p* = 7.45 × 10^−6^) and Integrable sequences are 0.71× less dense (*p* = 6.79 × 10^−4^).

Cumulative distribution function analysis confirms this inversion (Figure 6d) where Kolmogorov Smirnov tests versus Astral95 show significant distributional shifts for all categories with all *p <* 6.4 × 10^−5^. The systematic rightward shift of IDP and Knotted curves contrasts directly with the leftward shift of Random and Integrable sequences. The Kolmogorov Smirnov *D* statistic highlights these shifts with Random at 0.505 Integrable at 0.384 Anchor at 0.153 Fold-switching at 0.246 IDP at 0.313 and Knotted at 0.228.

This density inversion carries a direct statistical interpretation. Distinctive sequence-level features within IDPs knotted proteins and fold-switching proteins are strictly constrained by evolutionary selection [29,30] despite their physical diversity. The model compresses these shared grammatical signatures into compact manifold regions and produces high-density clusters. Random sequences lack coherent evolutionary grammar and consequently map to sparse peripheral regions. This density mapping demonstrates that functionally extreme proteins occupy highly clustered regions of the latent manifold and indicates implicit functional awareness despite the failure to achieve separable representations. The combination of high manifold density and negative Silhouette Scores (Figure 5) demonstrates that the model encodes a thermodynamic ensemble organized by sequence grammar rather than microscopic geometric identity (Figure S7). Physical conformational entropy undergoes a marked inversion in this latent space. Intrinsically disordered proteins possess high physical entropy and explore vast three-dimensional volumes but remain subject to rigorous evolutionary constraints to preserve flexibility. The language model captures this low evolutionary sequence entropy and packs these physically expansive proteins into the densest regions of the mathematical manifold.

### 2.6 Latent Space Exhibits Class-Invariant Geometric Turbulence

The coexistence of macroscopic separability between random and biological sequences alongside the microscopic conflation of topological phases delineates the operational boundaries of the latent representation. Evaluating the intrinsic continuous geometry of the manifold through persistent homology and gauge-field holonomy defects in the PCA-50D space uncovers the mathematical mechanism driving this macro-micro dissociation (Figure 7).

**Figure 7.**
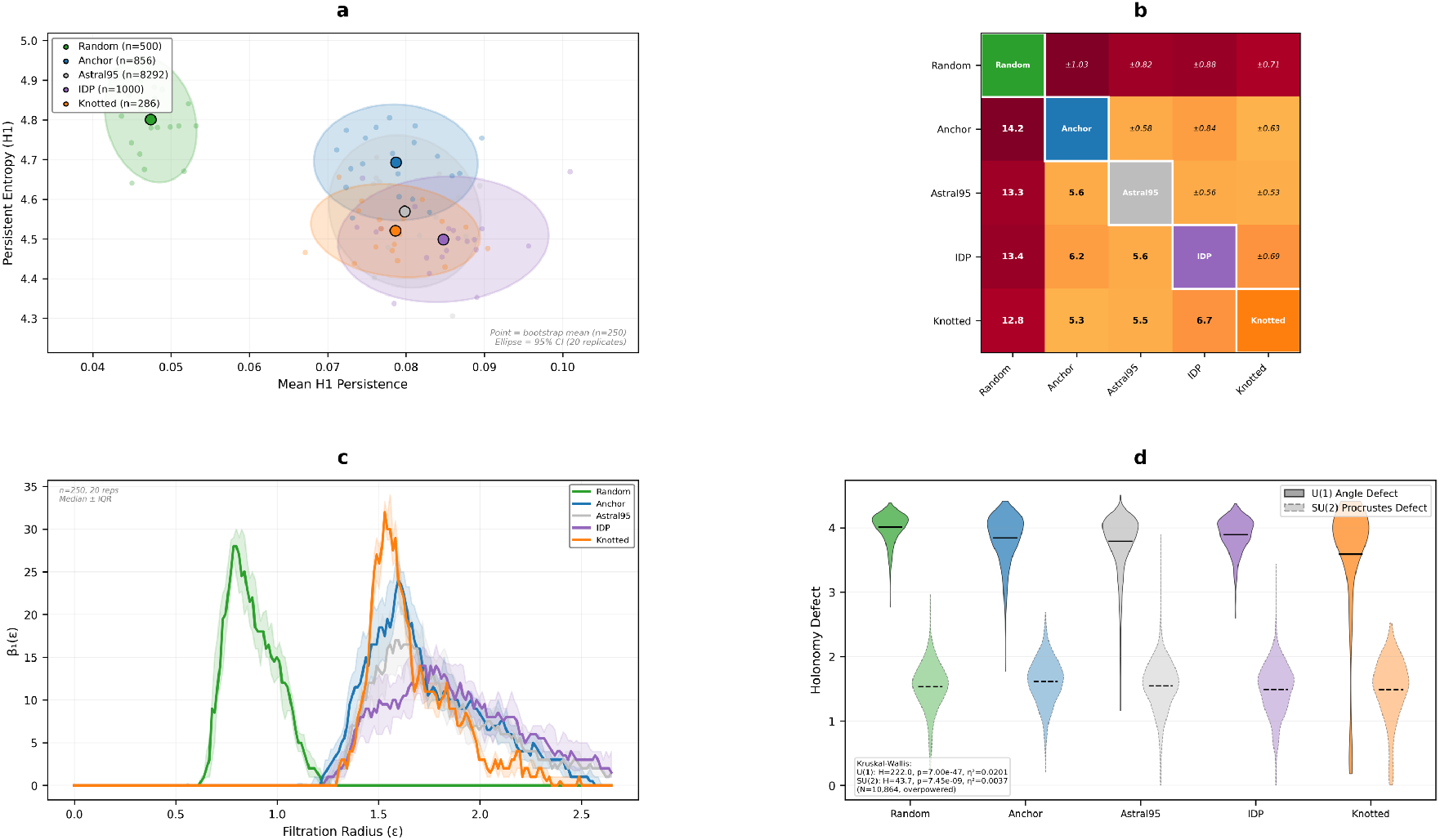
Persistent homology and gauge-field analysis of ESM-2 latent geometry. Five structural classes (Random *n* = 500, Anchor *n* = 856, Astral95 *n* = 8,292, IDP *n* = 1,000, Knotted *n* = 286) were analyzed in PCA-50D space. Fold-switching and Integrable categories were excluded to ensure equal sample sizes (*n* = 250, 20 replicates) for unbiased bootstrap resampling. **(a)** Topological complexity phase space mapping mean *H*_1_ persistence against persistent entropy. Ellipses denote 95% confidence regions. **(b)** Pairwise Wasserstein-2 distance matrix between *H*_1_ persistence diagrams. The lower triangle shows the mean distance, and the upper triangle shows the standard deviation across bootstrap replicates. **(c)** Betti-1 curves *β*_1_(*ε*) across filtration scales, displayed as median *±* interquartile range (IQR). **(d)** Holonomy defect distributions. Paired violin plots display the *U* (1) angle defect (solid, left half) and *SU* (2) Procrustes defect (dashed, right half) evaluated across local neighborhood loops for each class.

We restricted the topological analysis to five core categories including Random Anchor Astral95 IDP and Knotted proteins. We excluded the Fold-switching and Integrable categories because their sample sizes fall below the required bootstrap threshold of *n* = 250 to prevent confounding persistent entropy with sample-size artifacts.

Persistent homology reveals clear topological separation at the macroscopic level. Random sequences separate from all biological classes in the *H*_1_ phase space with a mean persistence of 0.047 and 95% CI [0.042, 0.053] (Figure 7a). This separation yields uniformly large Wasserstein-2 distances *W*_2_ *>* 12 (Figure 7b) while inter-biological distances remain moderate at *W*_2_ ≈ 5 to 7 (Kruskal-Wallis *H* = 54.72 *p* = 3.72 × 10^−11^). Betti-1 curves demonstrate that Random sequences peak earliest at *ε* ≈ 0.79 with *β*_1_ = 28.0 consistent with uniformly distributed short-lived loops (Figure 7c). Biological classes peak later between *ε* ≈ 1.5 and 1.7 with varying amplitudes that reflect structured persistent topological features (Figure S8). Anchor peaks at *β*_1_ = 24.0 Astral95 peaks at *β*_1_ = 17.0 IDP peaks at *β*_1_ = 14.0 and Knotted peaks at *β*_1_ = 32.0.

This structural pattern completely reverses at the microscopic level. Holonomy defect analysis measures the failure of parallel transport around local loops to provide a pointwise gauge-field diagnostic of local curvature (Figure 7d and Figure S9). The Kruskal-Wallis tests yield statistical significance with *H* = 222.0 (*p* = 7.00 × 10^−47^) for *U* (1) and *H* = 43.7 (*p* = 7.45*×*10^−9^) for *SU* (2) but the effect sizes remain negligible. Both 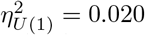 and 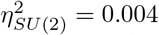 fall far below the 0.06 threshold across 10864 points. This demonstrates that local geometric curvature does not vary meaningfully across categories and establishes a state of class-invariant geometric turbulence.

This macro-micro dissociation resolves the apparent paradox of topological aliasing. The latent space contains macroscopic topological structure sufficient to distinguish random from biological sequences while its microscopic geometry remains uniformly turbulent and prevents the resolution of finer categorical distinctions. The model captures the global statistical regularities of protein sequence evolution but fails to learn the local geometric signatures that differentiate topological and thermodynamic phases. The macroscopic organization successfully identifies broad biological functional classes but querying the local neighborhood of a specific protein sequence yields predominantly geometric noise. This uniform local turbulence mathematically explains the inability of the continuous embedding geometry to yield precise conformational states or folding transition pathways.

### 2.7 Explicit Structural Encodings Partially Alleviate Topological Aliasing

The preceding analyses establish that ESM-2 conflates topologically distinct categories into overlapping latent regions. To determine whether this aliasing arises from the absence of explicit spatial information, we repeated the three class clustering analysis using SaProt 650M [43], a structure aware protein language model that encodes both amino acid sequences and Foldseek 3Di structural alphabet tokens [44] within a shared vocabulary. We evaluated 1250 proteins comprising 856 anchor, 261 knotted, and 133 fold-switching sequences. Intrinsically disordered proteins were excluded because they lack experimentally resolved static structures (Figure 8a).

**Figure 8.**
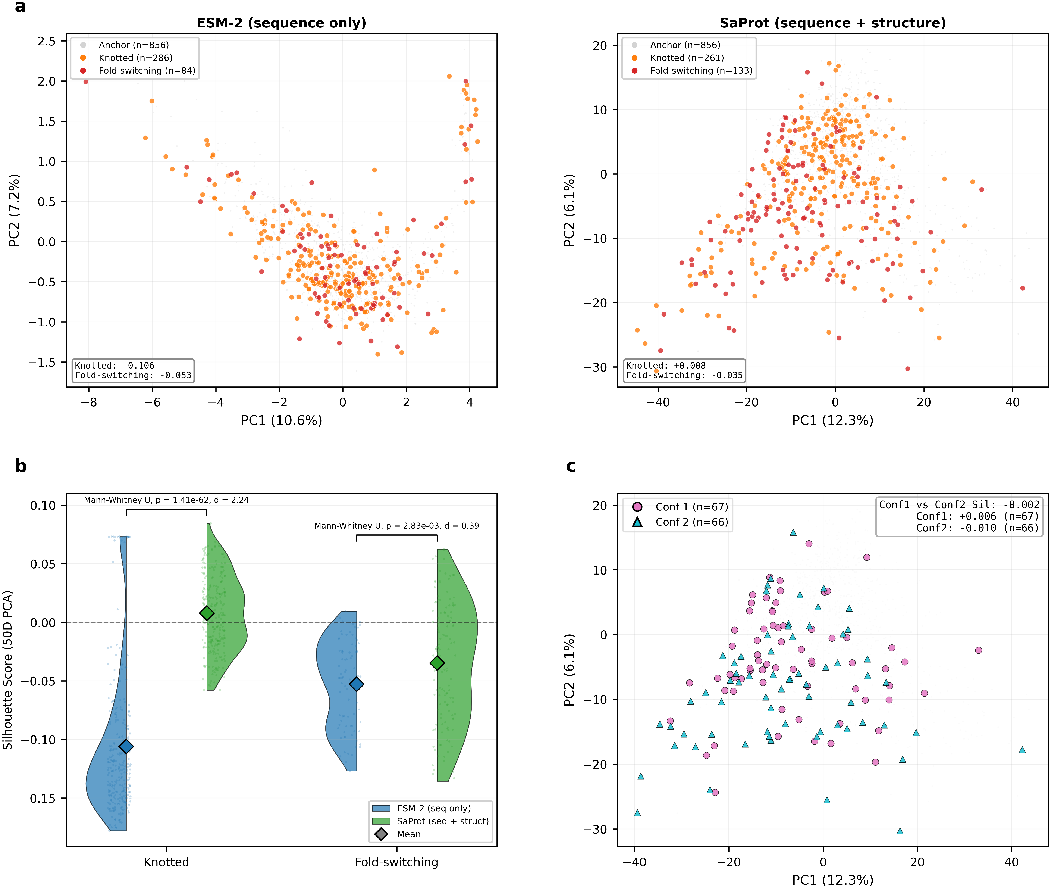
SaProt structure-aware control experiment. **(a)** Two dimensional PCA projections of protein embeddings from ESM-2 (sequence only, left) and SaProt (sequence and Foldseek 3Di structure tokens, right). Anchor proteins (gray, *n* = 856) form the background population. Knotted (orange) and fold-switching (red) proteins are overlaid. Inset boxes report mean Silhouette scores computed in the 50 dimensional PCA space. ESM-2 explained variance: PC1 = 10.6%, PC2 = 7.2%. SaProt explained variance: PC1 = 12.3%, PC2 = 6.1%. **(b)** Distribution of per sample Silhouette scores for knotted and fold-switching categories. Split violins compare ESM-2 (blue) and SaProt (green) embeddings. Diamonds mark category means and individual data points are jittered alongside. The dashed line marks the zero score boundary. The inclusion of explicit structural tokens increases the clustering score for knotted proteins while providing a smaller modification for fold-switching proteins. **(c)** Fold-switching proteins in the SaProt latent space colored by conformational state (conf1: circle, *n* = 67; conf2: triangle, *n* = 66). Alternative conformational states of the same protein map to overlapping latent regions, suggesting that static structural tokens may not fully resolve dynamic thermodynamic phases.

For knotted proteins, explicit structural input provided an observable reduction in topological aliasing. The mean Silhouette score increased from −0.106 (ESM-2) to +0.008 (SaProt), a shift confirmed by a one sided Mann-Whitney *U* test (*p* = 1.41 × 10^−62^, Cohen’s *d* = 2.24). The fraction of knotted proteins with positive Silhouette scores rose from 8 percent to 56 percent, indicating that more than half of the knotted sequences moved closer to their category centroid than to the anchor population (Figure 8b).

For fold-switching proteins, the improvement was marginal. The Silhouette score shifted from −0.053 to −0.035 (*p* = 2.83 × 10^−3^, Cohen’s *d* = 0.39), remaining negative and indicating persistent aliasing with the anchor population. Five nearest neighbor balanced accuracy across the three classes improved from 0.465 (ESM-2) to 0.597 (SaProt), exceeding the chance level of 0.333 but remaining below reliable separability.

To directly test whether structure aware encoding can distinguish alternative fold states, we embedded both conformational partners for 67 fold-switching protein pairs and computed the Silhouette score separating conf1 from conf2 labels. The resulting score of −0.002 indicates high latent overlap between the two fold states (Figure 8c). Paired conformations of the same protein are closer to each other than random fold-switching pairs in 50 dimensional PCA space (42.74 *±* 13.16 *±* versus 46.75 *±* 12.46; *t* = −2.52, *p* = 0.012), confirming partial preservation of protein level identity. However, this proximity does not translate into categorical discriminability. The model maps conformational partners to the same latent neighborhood despite their distinct three dimensional folds.

These results delineate a boundary in the representational capacity of structure-aware sequence models. Explicit structural tokens alleviate aliasing for static topological anomalies whose geometric signatures persist across crystallographic snapshots. For dynamic topological phases governed by thermodynamic equilibria between alternative folds, the aliasing persists because the sequence-centric architecture lacks the capacity to represent environment-dependent conformational selection.

## 3 Discussion and Conclusions

The findings presented in this study clarify the representational capacity of purely data driven sequence modeling. The consistently negative Silhouette Scores for intrinsically disordered proteins knotted proteins and fold-switching proteins demonstrate that the model conflates physically distinct categories that share overlapping sequence statistics. Although this evolutionary grammar effectively encodes functional interaction networks and predicts phase separation motifs [16, 18, 19], it fails to resolve exact spatial geometry. For standard proteins satisfying the Anfinsen paradigm the evolutionary sequence statistics and the physical folding rules are strongly correlated and allow the model to serve as an effective structural proxy. This correlation diverges for dynamic proteins and topologies governed by complex thermodynamic phase behavior, where conformational states are heavily modulated by transient structural elements and chemical environments [17, 20, 26, 27]. Resolving such context dependent mappings highlights the growing necessity for architectures that can dynamically adjust protein representations based on specific cellular and tissue environments [45]. Our region-replacement experiment (Figure 5b) rules out the sequence dilution hypothesis and establishes that this topological aliasing is an intrinsic characteristic of the representational framework itself. The holonomy defect analysis (Figure 7d) reveals the microscopic mechanism behind this representational limit. Local geometric curvature in the latent space is class invariant (*η*^2^ *<* 0.06) and produces a uniform geometric turbulence. The latent manifold is macroscopically distinguishable but microscopically indistinguishable. While advances in topological computation can extract highly localized cycle representatives in physical biological data [46], the latent space lacks this fine-grained structural identifiability. While recent discrete geometric frameworks successfully map real-space structural phase transitions and thermodynamic phase behavior using spectral entropy and integrability residuals [36, 37], the uniform turbulence of the latent space precludes such microscopic resolution. This mathematically characterizes the topological boundary of the model when evaluating proteins that undergo dynamic conformational remodeling and structural phase transitions.

While these findings define the representational boundaries of current protein language models the analytical methodology contains specific constraints. The application of UMAP for density estimation introduces a dependence on hyperparameters. This step remains mathematically necessary to circumvent the curse of dimensionality in the fifty-dimensional principal component space as detailed in Figures S5 and S6. Furthermore the Hasimoto integrability error *E*[*n*] relies on static crystal structures. Static models may not fully capture the continuous conformational flexibility inherent to IDPs and fold-switching proteins. Extending this differential geometric analysis to ensemble structures derived from nuclear magnetic resonance or molecular dynamics simulations will provide a more comprehensive dynamic evaluation. The region-replacement experiment provides rigorous statistical control for the 84 fold-switching proteins analyzed here. Expanding this analysis to larger fold-switching datasets as they become experimentally available will further validate the generality of these representational observations.

Recognizing these representational boundaries informs the developmental trajectory for biomolecular representation learning. Current sequence derived embeddings are frequently utilized to predict complex dynamic phenomena including allosteric regulation and phase separation. Task specific supervised fine tuning of these language models via accessible community platforms like SaprotHub can partially bridge performance gaps for localized functional and fitness predictions [47, 48]. The demonstration of topological aliasing suggests that these applications operate in a representational space that fundamentally lacks the geometric resolution to distinguish thermodynamic phases. Our SaProt control experiment (Figure 8) corroborates this assessment: appending static Foldseek 3Di structural tokens to the input vocabulary substantially reduces aliasing for knotted proteins (Silhouette −0.106 → +0.008, Cohen’s *d* = 2.24) yet leaves fold-switching conf1 and conf2 states indistinguishable (Silhouette = −0.002). The improvement is therefore confined to static topological anomalies whose geometric signatures are already captured by a single crystallographic snapshot, while multi-state thermodynamic phases remain aliased. This result indicates that the representational bottleneck lies not in the absence of structural input per se, but in the lack of conformational dynamics encoding.

Future models require the integration of physical laws governing topological and thermodynamic phases rather than relying exclusively on sequence statistics. Continued scaling of model parameters and sequence data will further refine evolutionary modeling but capturing genuine physical causality necessitates architectures that extend beyond sequence prediction. Recent multi-modal foundation models that explicitly align sequence, structure, and binding-site representations offer an initial step in this direction [49]. The development of conformational dynamics world models offers a necessary advancement. Such frameworks learn the transition rules of the physical energy landscape and the cooperative kinetics of barrier crossing [50] rather than merely mapping static sequence distributions. This approach imposes temporal and physical constraints on the latent space to accurately represent allosteric networks and folding trajectories. Incorporating physics informed loss functions that penalize violations of geometric invariants during pre-training can force the model to preserve microscopic geometric information. Indeed, recent advancements in artificial protein evolution demonstrate that explicitly integrating both structural constraints and evolutionary language models substantially increases engineering success rates compared to purely sequence-based approaches [51].

As recently demonstrated by Frank et al. [52], capturing genuine physical correlations in Euclidean space necessitates architectures equipped with explicit mechanisms that rigorously respect three-dimensional translational and rotational symmetries. The absence of such explicit equivariant encodings in purely sequence based models mathematically explains the geometric turbulence and topological blindness observed in our analysis of ESM-2. Utilizing equivariant neural architectures therefore provides a necessary mechanism to retain the local spatial features that mean-pooling currently discards. The topological aliasing documented in this study characterizes a fundamental boundary of purely data driven protein representations. Advancing beyond this limitation requires fusing the statistical capacity of large language models with the geometric precision and physical constraints of thermodynamic principles.

## 4 Methods

### 4.1 Dataset Construction

We constructed a dataset of 11,068 protein sequences spanning seven functional categories to probe the latent space of ESM-2 Figure S1. The structural reference set comprises 8,292 sequences from the ASTRAL SCOP 95 percent non-redundant set [42, 53–55] selected via the PISCES server [56, 57]. Selection criteria required X-ray crystallography resolution ≤ 2.0 Å, *R*-factor ≤ 0.2, sequence identity ≤ 25 percent, and chain lengths between 40 and 300 residues. This set covers all four major SCOP classes. We included 856 high-fidelity reference proteins from the Protein Data Bank, previously curated via the PISCES server [34] using identical structural criteria while strictly excluding chains with breaks or disorder. These physical structures serve as fixed reference points in the embedding manifold and provide the absolute ground truth for computing the microscopic geometric integrability error *E*[*n*]. The intrinsically disordered proteins category contains 1,000 sequences from the DisProt database [12] experimentally validated to lack stable tertiary structure under physiological conditions. The knotted proteins category includes 286 sequences derived from the KnotProt database [28] with experimentally or computationally confirmed topological knots in their backbone. The fold-switching proteins category consists of 84 literature curated sequences [21] experimentally demonstrated to adopt multiple distinct stable folds in response to cellular cues. We incorporated 50 integrable helices sequences exhibiting near perfect integrability under the Hasimoto map. The random control sequences comprise 500 sequences generated by sampling from the UniProt [58] derived natural amino acid frequency distribution with lengths drawn from a normal distribution matching empirical protein length statistics.

### 4.2 ESM-2 Embedding Extraction

Sequence embeddings were extracted using the ESM-2 3B model from Hugging Face Transformers [59] under the identifier facebook/esm2 t36 3B UR50D. Sequences were truncated to a maximum length of 1,024 tokens. Sequence level embeddings of 2,560 dimensions were obtained by attention masked mean pooling of the last hidden state while strictly excluding padding tokens. The embedding vector **e** is defined as Σ_*i*_ *m*_*i*_**h**_*i*_*/* Σ_*i*_ *m*_*i*_ where **h**_*i*_ represents the hidden state at position *i* and *m*_*i*_ represents the attention mask. Inference was performed using mixed precision FP16 on an NVIDIA A40 GPU with a batch size of 8. All random seeds were fixed to 42 to ensure computational reproducibility.

### 4.3 Dimensionality Reduction

Embeddings were standardized to zero mean and unit variance per dimension using StandardScaler [60] before principal component analysis. The PCA operation reduced the standardized 2,560 dimensional embeddings to 50 dimensions to serve as the primary analysis coordinate system. The 50 dimensional PCA space robustly preserves global pairwise distance structure and serves as the foundation for all geometric analyses including Silhouette Scores, Ollivier-Ricci curvature, persistent homology, and holonomy defects.

UMAP [61] was applied to the 50 dimensional PCA embeddings and projected to two dimensions to facilitate kernel density estimation. The UMAP parameters included 15 nearest neighbors, a minimum distance of 0.1, the Euclidean metric, and a random state of 42. The emphasis of UMAP on local structure proves advantageous for density mapping Figure S3. The rationale for this dual approach is mathematically detailed in Figures S5 and S6. High dimensional kernel density estimation fails because kernel weights collapse to near zero under the curse of dimensionality. Conversely, 50 dimensional topological computation via persistent homology remains completely valid because it relies on distance rankings rather than absolute kernel weights. A t-SNE projection [62] was computed for comparison using a perplexity value of 30.

### 4.4 Computation of Microscopic Hasimoto Integrability Error

We computed the Hasimoto integrability error *E*[*n*] for 9,195 structurally resolved sequences to mathematically quantify the microscopic geometric variation of the protein backbone in physical space. Although derived from continuous gauge field theory and the nonlinear Schrödinger equation, *E*[*n*] functions strictly as a discrete local order parameter for backbone symmetry.

Given a discrete protein backbone represented by *C*_*α*_ coordinates 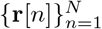, we constructed the discrete Frenet frame to obtain the bond angle *κ*[*n*] and torsion angle *τ* [*n*] at each residue *n*. The discrete Hasimoto transform maps the three dimensional geometry onto a complex scalar field *ψ*[*n*] = *κ*[*n*] exp(*i* Σ *τ* [*k*]). The real part of the exact effective potential *V*_re_[*n*] is derived algebraically from local curvature ratios and torsion angles according to the following equation.

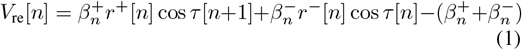

The terms *r*^*±*^[*n*] = *κ*[*n ±* 1]*/κ*[*n*] represent local curvature ratios. The terms 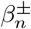 denote sequence dependent bond stiffness parameters which we approximated as a uniform coupling *β* in this study. The integrability error is defined as the absolute residual of the discrete nonlinear Schrödinger dispersion relation.

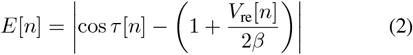

Geometrically *E*[*n*] measures the exact degree of discrete helical symmetry breaking. A value approaching zero corresponds to geometrically regular and integrable *α*-helices. A large positive value indicates broken symmetry regions characterized by high geometric variation such as unstructured loops and *β*-strands. Rigorous mathematical derivations establishing this exact discrete decomposition as a kinematic identity rather than a dynamical governing equation are provided by Wang [34]. Recent applications of this framework further validate *E*[*n*] as a quantitative tool for identifying piecewise integrable helical islands [35], extracting sub-residue geometric transitions via spectral entropy [36], and mathematically characterizing biological order [37].

### 4.5 Geometric Analysis

Ollivier-Ricci discrete curvature was computed in the 50-dimensional PCA space on a *k*-nearest neighbor graph (*k* = 20, Euclidean metric). For each edge (*x, y*), the curvature was estimated as *κ*(*x, y*) = 1 − *W* (*µ*_*x*_, *µ*_*y*_)*/d*(*x, y*), where *W* is the Wasserstein distance approximated by the mean of minimum matching distances between the *k*-neighborhoods of *x* and *y*. The per-point curvature was obtained by averaging edge curvatures over all *k* = 20 neighbors. Local dimension and condition numbers were computed from local covariance matrices of *k*-neighborhoods using eigenvalue decomposition in 50D PCA space.

### 4.6 Silhouette Analysis and Region Replacement

Per-sample Silhouette Scores were computed in PCA 50D space using Euclidean distances and the 7-class label system (Random, Integrable, Anchor, Astral95, Fold-switching, IDP, Knotted). For the region-replacement experiment, distinctive regions of three extreme categories were replaced: fold-switching interface units (IFU regions) for Fold-switching, disordered regions for IDPs, and knotted regions for Knotted proteins. Replacement sequences were drawn from matched ASTRAL95 proteins. Both full-length and region-replaced sequence embeddings were extracted from ESM-2, independently projected to PCA 50D, and Silhouette Scores recomputed to assess whether distinctive regions contribute to category separability.

### 4.7 Density Estimation

Manifold density was computed using kernel density estimation (KDE) in 2D UMAP space rather than the original high-dimensional embedding space. High-dimensional KDE suffers from the curse of dimensionality (Figures S5 and S6): in 50D PCA space, the Gaussian kernel weight at the mean pairwise distance collapses to approximately 2.85 × 10^−3^ under Scott’s bandwidth, rendering density estimates unreliable. In contrast, 2D UMAP density provides a continuous distribution suitable for statistical analysis. We used scikit-learn [60] `KernelDensity` with Gaussian kernel and bandwidth determined by Scott’s rule, 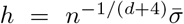. Fold-change values were computed relative to the Astral95 reference set, with significance assessed by Mann-Whitney *U* tests and Kolmogorov–Smirnov tests.

### 4.8 Holonomy Defect and Persistent Homology

Persistent homology was computed using Ripser [63] (maxdim = 2) on bootstrap-resampled subsets (*n* = 250, 20 replicates per category) in PCA 50D space with Vietoris-Rips filtration. For each replicate, we computed *H*_1_ persistence diagrams, from which persistent entropy and mean persistence were derived. Pairwise Wasserstein-2 distances between *H*_1_ diagrams were computed across all category pairs. Fold-switching (*n* = 84) and Integrable (*n* = 50) were excluded because their sample sizes fall below the bootstrap threshold of *n* = 250.

To measure the intrinsic microscopic curvature of the ESM-2 latent space, we computed holonomy defects utilizing a discrete graph-based gauge field formalism. For each data point in the PCA 50D space, we identified a minimal closed loop *γ* = {*x*_1_, *x*_2_, …, *x*_*m*_, *x*_1_} within its *k*-nearest neighbor graph (*k* = 20). The parallel transport of a tangent vector around this loop is governed by the discrete Wilson loop operator *W* (*γ*), adapting the lattice gauge theory formalism [64] to the data manifold:

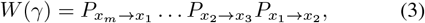

where *P*_*x*→*y*_ is the orthogonal projection matrix aligning the local tangent spaces (computed via local PCA neighborhoods) of adjacent nodes, a discrete parallel transport construction central to the analysis of vector fields on data manifolds [65]. We extracted two geometric invariants from the returned vector: the *U* (1) angular defect, defined as the accumulated phase angle mismatch (analogous to the quantal geometric phase accumulated along cyclic parameter evolutions [66]), and the *SU* (2) Procrustes defect, quantifying the rotational matrix distance from the identity. Effect sizes (*η*^2^) from Kruskal-Wallis tests assessed whether these defect distributions differ meaningfully across categories.

In the context of language model representations, a low holonomy defect indicates a locally flat representational manifold where sequence transitions are geometrically smooth. Conversely, a consistently high holonomy defect reflects a state of “geometric turbulence”—the latent space is tightly folded at the microscopic level, mechanically explaining the topological aliasing observed between structurally distinct proteins.

### 4.9 Statistical Analysis

Spearman rank correlations, Mann-Whitney *U* tests, Kolmogorov–Smirnov tests, and Kruskal-Wallis tests were computed using SciPy [67]. Effect sizes include Cohen’s *d, η*^2^, and Kolmogorov–Smirnov *D* statistic. Significance threshold was set at *p <* 0.05 for individual tests, with Bonferroni correction applied for multiple testing where indicated. All tests are two-tailed unless otherwise specified.

### 4.10 SaProt Structure-Aware Control

To assess whether explicit structural encoding alleviates topological aliasing, we extracted embeddings from SaProt 650M [43], a protein language model trained on interleaved amino acid and Foldseek 3Di structural alphabet tokens [44]. Three-dimensional coordinates for each protein were processed through Foldseek to obtain per-residue 3Di tokens, which were interleaved with standard amino acid characters to form SaProt input sequences. The dataset comprised 1,250 proteins (856 anchor, 261 knotted, 133 fold-switching including both conformational partners for 67 pairs); intrinsically disordered proteins were excluded because they lack experimentally resolved static structures. Sequence-level embeddings (1,280 dimensions) were obtained by mean pooling the last hidden state. Dimensionality reduction and Silhouette Score computation followed the procedures described in the preceding subsections, with the label system reduced to three classes (anchor, knotted, fold-switching). ESM-2 versus SaProt distributional shifts were tested with one-sided Mann-Whitney *U* tests (alternative: SaProt *>* ESM-2) and effect sizes reported as Cohen’s *d*. Three-class discriminability was assessed by 5-nearest-neighbor balanced accuracy under 5-fold stratified cross-validation. Conformational partner analysis computed pairwise Euclidean distances in PCA 50D space between conf1 and conf2 embeddings of the same fold-switching protein and compared them to distances between random fold-switching pairs using a two-sample *t*-test.

## Acknowledgments

This work was supported in part by the Computing and Data Center of Xinjiang University. We acknowledge the computing resources and technical support provided by the Computing and Data Center of Xinjiang University.

## Data and code availability

The complete analytical pipeline for this study is freely available at https://github.com/wyqmath/ESM-Latent-Topology. Raw sequence and structural data were obtained from the ASTRAL SCOPe database [54, 55] (https://scop.berkeley.edu/) and the RCSB Protein Data Bank (PDB, https://www.rcsb.org/).

## Supplementary Information

### Rationale for 2D UMAP Density Over 50D PCA Density

High-dimensional kernel density estimation (KDE) suffers from the curse of dimensionality (Figures S5 and S6). In 50D PCA space, the mean pairwise Euclidean distance between points follows the scaling 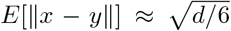; at *d* = 50, this yields a distance of approximately 2.89. With Scott’s rule bandwidth *h* = *n*^−1*/*(*d*+4)^*σ* where *n* = 11,068 and *d* = 50, we obtain *h* ≈ 0.84. The Gaussian kernel *K*(*x*) = exp(−∥*x*∥^2^*/*2*h*^2^) evaluated at the mean distance yields a weight of approximately 2.85 × 10^−3^, so most samples contribute near-zero local density mass.

This causes the density distribution to collapse into discrete, quantized values. Controlled simulations (Figure S6) confirm that 50D KDE with *n* = 1,000 Gaussian samples produces only 1 unique density value at 10-decimal rounding, whereas 2D KDE yields 1,000 unique values, a 1,000× resolution difference. The required sample size for reliable 50D KDE scales as approximately *k*^*d*^ (where *k* is the resolution per dimension), requiring ∼ 8.88 × 10^34^ samples versus our ∼ 10^4^, a deficit of ∼ 10^30^.

In contrast, 2D UMAP density provides a continuous distribution suitable for statistical analysis. The choice of UMAP 2D for density estimation and PCA 50D for geometric/topological analyses reflects the complementary strengths of each space: UMAP preserves local neighborhood structure needed for density estimation, while PCA preserves global distance structure needed for Silhouette Scores, persistent homology, and curvature analysis.

### Thermodynamic Interpretation of Density Inversion

The density inversion (Figure 6), in which IDPs (1.36*×*), knotted proteins (1.24*×*), and fold-switching proteins (1.19*×*) occupy denser manifold regions than the Astral95 structural baseline, reveals the distinction between physical conformational space and learned sequence-grammar space. In physical conformational space, the density of states satisfies *ρ* ∝ exp( −*S/k*_*B*_), where high conformational entropy (disorder) corresponds to low density. This physical disorder can be rigorously quantified through discrete geometric invariants such as spectral entropy and integrability residuals, which mathematically characterize the biological structural order of the backbone [36, 37]. In the ESM-2 latent space, this relationship is inverted because the model organizes sequences by evolutionary sequence grammar rather than by these precise physical properties.

IDPs, despite their conformational heterogeneity, share distinctive sequence-level features (enrichment in polar and charged residues, low-complexity regions, and specific amino acid biases) that are strongly constrained by evolutionary selection. The model compresses these shared grammatical signatures into compact manifold regions, creating high-density clusters. Random sequences, which lack coherent evolutionary grammar, are mapped to sparse peripheral regions (0.82× baseline density).

This can be formalized as *ρ*_latent_ ∝ exp( −*S*_grammar_*/k*_*B*_), where *S*_grammar_ is the *grammatical entropy*, the diversity of sequence-level evolutionary signatures, rather than physical conformational entropy. Categories with distinctive, constrained grammar (IDPs, knotted proteins) have low grammatical entropy and high latent density, while categories with diverse or absent grammar (random, integrable) have high grammatical entropy and low latent density (Figure S7).

### Evaluating the Dilution Hypothesis

The dilution hypothesis posits that ESM-2’s inability to separate topologically distinct proteins arises because mean-pooling dilutes the signal from distinctive regions (e.g., knotted cores, disordered segments, fold-switching interface units) with embedding vectors from non-distinctive flanking domains. If true, replacing the distinctive regions with matched ASTRAL95 sequences should substantially change the Silhouette Scores.

We tested this by constructing region-replaced embeddings for three extreme categories: Fold-switching (IFU regions replaced), IDP (disordered regions replaced), and Knotted (knotted regions replaced). Both full-length and region-replaced embeddings were independently projected to PCA 50D, and per-sample Silhouette Scores were recomputed with the 7-class label system. The near-zero changes (Fold-switching: +0.1%; IDP: +0.6%; Knotted: +0.0%) effectively rule out the dilution hypothesis, establishing that topological aliasing is an intrinsic limitation of the model rather than a consequence of information dilution. Single-sample tracking analysis (Figure S4) confirms that the per-sample perturbations are uniformly negligible.

**Figure S1.**
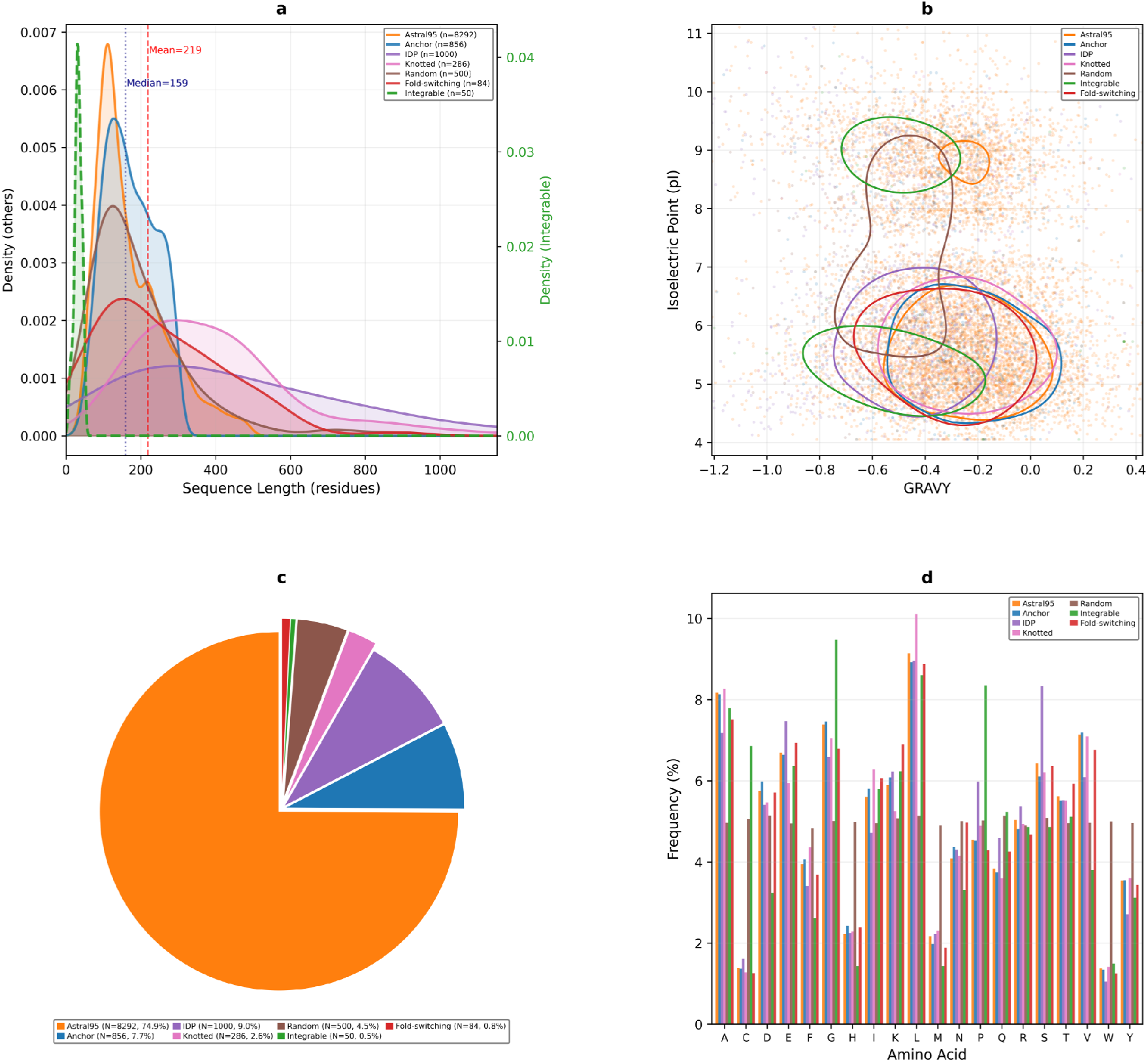
Dataset-wide statistical overview. Baseline statistics and physicochemical profiling for 11,068 protein sequences across seven categories (Astral95, Anchor, IDP, Knotted, Random, Integrable, Fold-switching). **(a)** Kernel density estimates (KDE) of sequence length. Across all sequences, length is 218.9 *±* 240.1 aa (mean *±* s.d.), median 159 aa, range [10, 7073] aa; the primary display window is clipped to the 1st–99th percentile [47, 1152] aa to suppress extreme long-tail effects. The Integrable subset (*n* = 50) is plotted on a separate right *y*-axis to preserve visibility. **(b)** Two-dimensional pI–GRAVY landscape (*n* = 11,032 valid sequences), colored by the seven categories with class-specific 2D KDE contours. GRAVY spans [ −2.765, 2.027] (mean −0.309 *±* 0.301), and pI spans [4.050, 12.000] (mean 6.602 *±* 1.704). The global Spearman correlation is *ρ*(GRAVY, pI) = −0.146 (*p* = 1.95 × 10^−53^), indicating a weak negative association. **(c)** Pie chart of category composition: Astral95 74.9% (*n* = 8,292), IDP 9.0% (*n* = 1,000), Anchor 7.7% (*n* = 856), Random 4.5% (*n* = 500), Knotted 2.6% (*n* = 286), Fold-switching 0.8% (*n* = 84), Integrable 0.5% (*n* = 50). **(d)** Grouped bar plot of amino-acid composition by category (20 standard amino acids). After sequence cleaning, the dataset contains 2,413,527 residues; the global top-5 amino acids are L (8.96%), A (7.79%), G (7.08%), S (6.80%), and V (6.79%). Cleaning rules match the main text: X removed, U→C, B→N, Z→Q.

**Figure S2.**
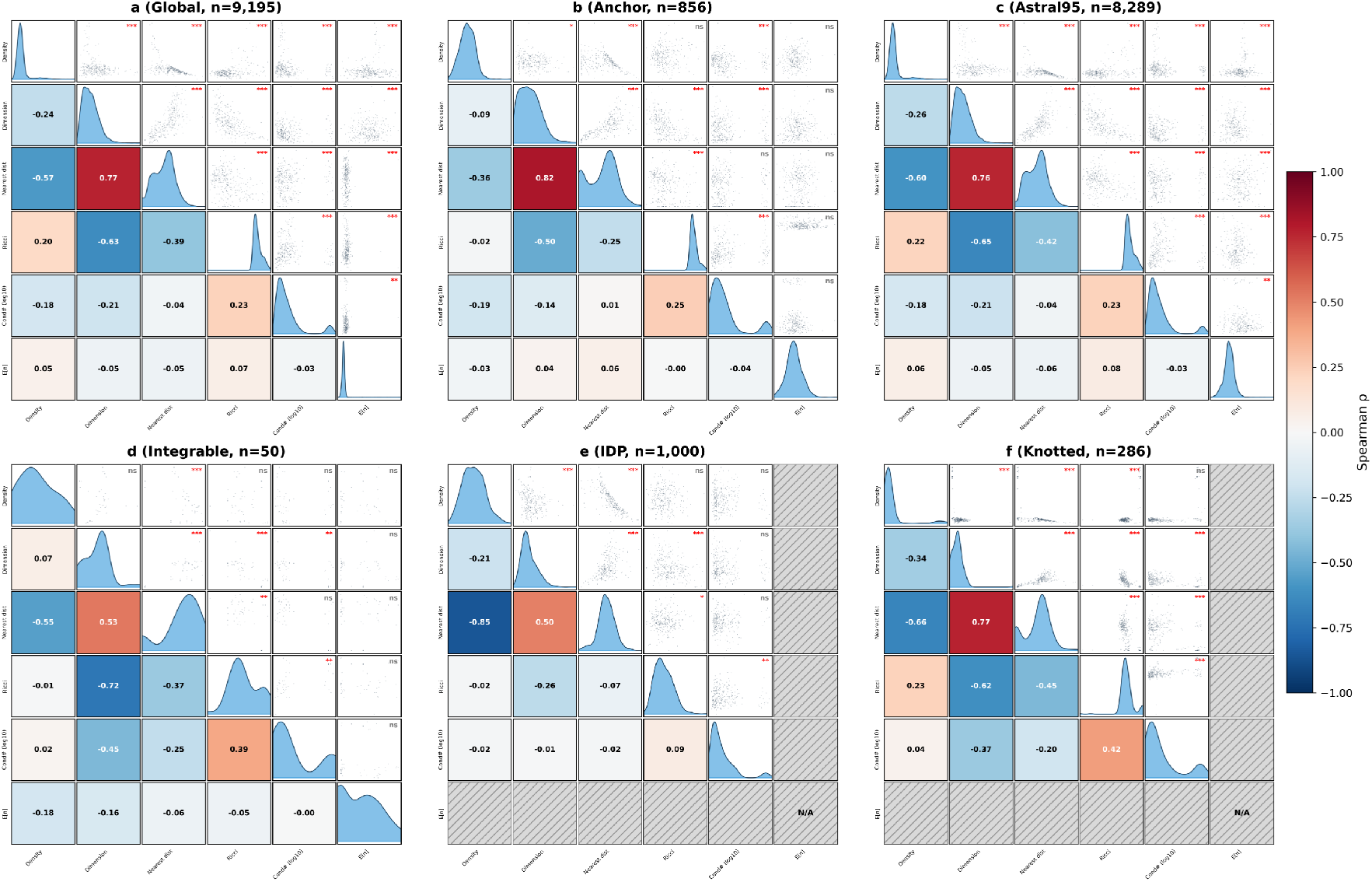
Category-decoupled correlation analysis. To control for category confounding, Spearman rank correlation matrices of six local geometric variables (Local Density, Local Dimension, Nearest Distance, Ricci Curvature, Condition Number (log_10_), *E*[*n*]) were computed separately on the global dataset and per-category subsets, displayed in a pair-grid format (diagonal: KDE density; lower triangle: correlation coefficient heatmap; upper triangle: scatter + significance marker). Geometric variables were computed from PCA-50D space using *k* = 20 nearest neighbors. **(a)** Global (*n* = 9,195, samples with *E*[*n*]): the highest correlation is observed between Local Dimension and Nearest Distance (*ρ* = +0.767); *E*[*n*] shows negligible correlations with all geometric variables (max |*ρ*(*E*[*n*], ·)| = 0.067, Ricci Curvature, *p* = 1.27 × 10^−10^). **(b)** Anchor (*n* = 856): this primary correlation remains consistent (*ρ* = +0.818); all correlations involving *E*[*n*] are non-significant (max |*ρ*| = 0.055, *p* = 0.107). **(c)** Astral95 (*n* = 8,289): the correlation structure closely mirrors the global pattern, peaking at *ρ* = +0.763 for the same variable pair; *E*[*n*] versus Ricci Curvature yields *ρ* = +0.079 (*p* = 4.86 × 10^−13^). **(d)** Integrable (*n* = 50): the small sample size alters the structural pattern, such that Local Dimension and Ricci Curvature exhibit the highest correlation (*ρ* = −0.720); all correlations involving *E*[*n*] are non-significant (max |*ρ*| = 0.181, *p* = 0.208). **(e)** IDP (*n* = 1,000) and **(f)** Knotted (*n* = 286): *E*[*n*] rows/columns are masked (grey hatching, N/A) because these categories lack a structurally meaningful *E*[*n*] definition. For IDPs, the highest correlation occurs between Local Density and Nearest Distance (*ρ* = −0.855). For Knotted proteins, Local Dimension and Nearest Distance show the maximum correlation (*ρ* = +0.766). Regarding cross-panel consistency, the pairs Local Density versus Nearest Distance, Local Dimension versus Nearest Distance, and Local Dimension versus Ricci Curvature recur in the top-3 |*ρ*| pairs across all six panels, confirming that these geometric couplings are category-invariant manifold structural invariants.

**Figure S3.**
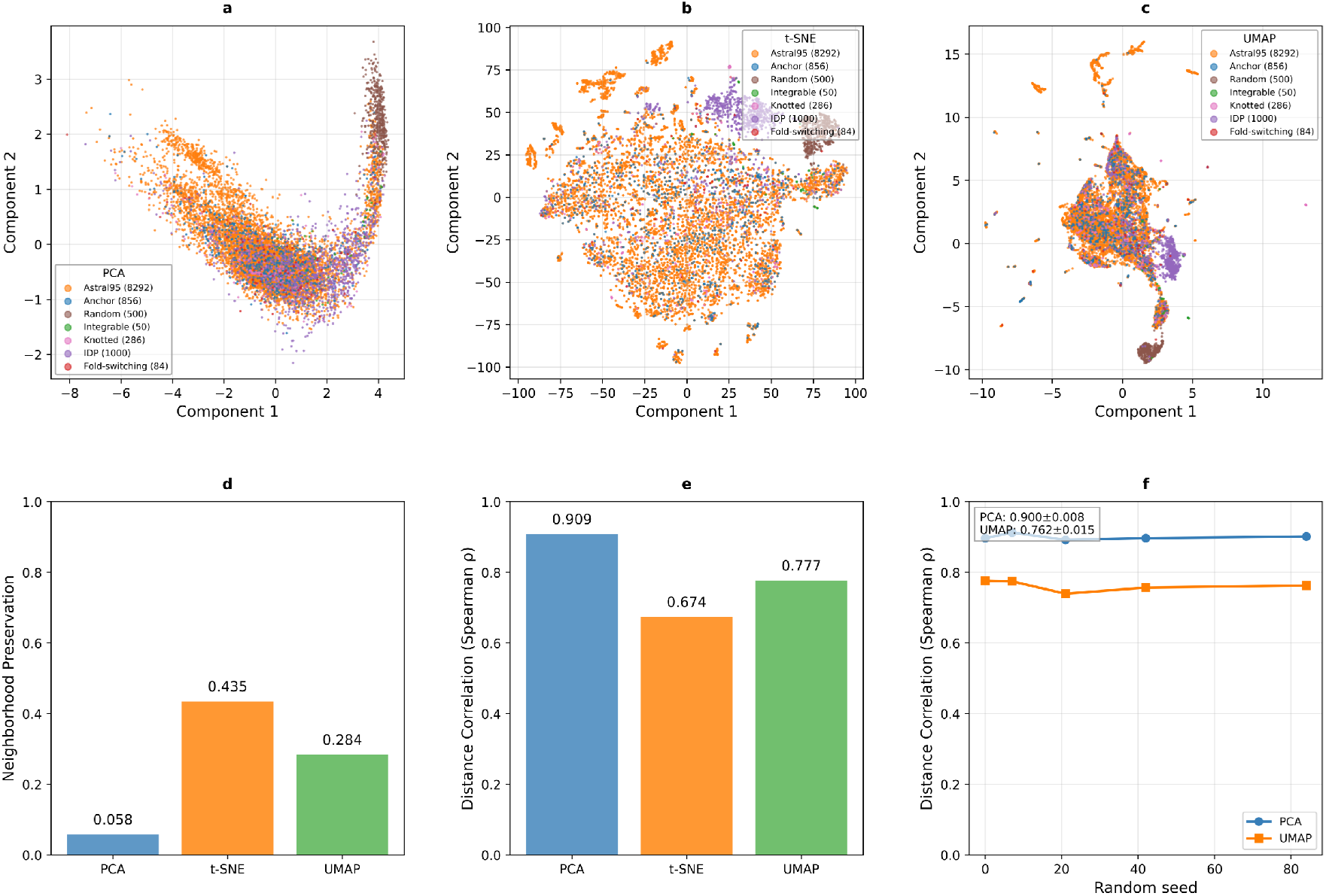
Robustness analysis of dimensionality reduction methods supporting the PCA choice in Figure 3. Using 11,068 ESM-2 sequence embeddings (first reduced to PCA 50D), we compared PCA, t-SNE, and UMAP in terms of manifold fidelity and robustness. **(a–c)** 2D scatter projections (category-colored) visualize their geometric organization differences. **(d)** Neighborhood Preservation (*k* = 15): t-SNE = 0.435, UMAP = 0.284, PCA = 0.058, indicating stronger local-neighborhood retention for t-SNE/UMAP. **(e)** Global distance fidelity (Spearman distance correlation to 50D pairwise distances): PCA = 0.909, UMAP = 0.777, t-SNE = 0.674, showing that PCA best preserves global geometry. We further quantified category-level local clustering using kNN purity (*k* = 15) to validate visual impressions: for IDP, PCA = 0.2703, t-SNE = 0.7180, UMAP = 0.6797; and in all 7 categories, the highest local purity was achieved by t-SNE, confirming that t-SNE/UMAP more strongly emphasize local class compaction. **(f)** Cross-seed robustness comparison (seed ∈ {0, 7, 21, 42, 84} ) plotting distance-correlation trajectories for PCA and UMAP with mean *±* s.d. annotation. PCA achieves DC = 0.9001 *±* 0.0079, while UMAP achieves DC = 0.7621 *±* 0.0150; UMAP variability is 1.91× higher (std ratio), and PCA has a +0.1379 mean global-fidelity advantage. Together, these results indicate that although t-SNE/UMAP better emphasize local neighborhoods, PCA simultaneously provides superior global distance preservation and better cross-seed reproducibility, justifying the PCA coordinate system used in main-text Figure 3. All correlations are two-tailed Spearman tests; UMAP parameters were fixed at *n*_neighbors = 15 and min_dist = 0.1. See Table S1 for complete quantitative summary.

**Figure S4.**
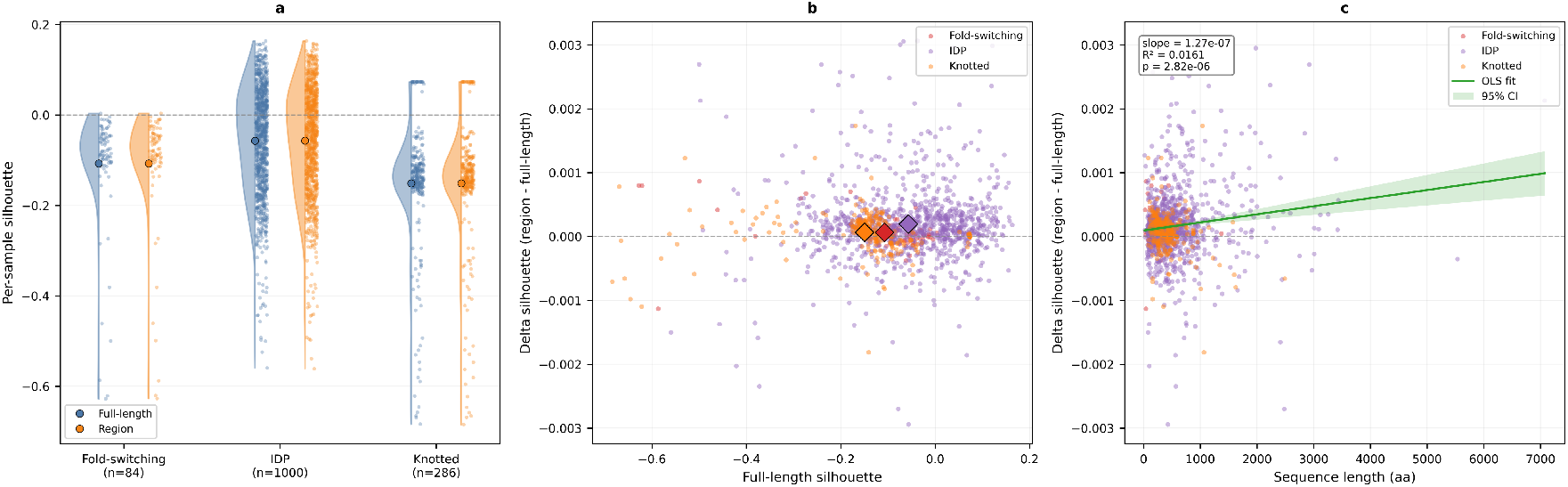
Single-sample micro-tracking of topological aliasing (supporting Figure 5). Using full-length and region-replaced embeddings for all 11,068 proteins, we quantified sample-level silhouette changes for three extreme categories (Fold-switching, IDP, Knotted). All silhouette scores were computed in PCA-50D space using the 7-class label system (Random, Integrable, Anchor, Astral95, Fold-switching, IDP, Knotted). **(a)** Raincloud distributions (left panel) show per-sample silhouettes for full-length vs region-replaced embeddings in each extreme category (half-violin + jittered points), enabling direct distribution-level comparison. **(b)** DeltaSilhouette scatter (middle panel): *x* = full-length silhouette, *y* = Δsilhouette = sil_region_ − sil_full-length_, colored by category, with category means highlighted by diamond markers; the horizontal dashed line indicates *y* = 0. To prevent visual scaling from being dominated by rare outliers, panels (b)(c) apply a unified 99th-percentile filter on |Δsilhouette| across all extreme samples (*q*_99_ = 0.003068), retaining 1356/1370 samples for plotting. **(c)** Length-attribution panel (right): *x* = sequence length (aa), *y* = Δsilhouette, with a green OLS fit line and a shaded 95% confidence interval. Filtered-fit statistics: slope = +1.266300 × 10^−7^, *R*^2^ = 0.016075, *p* = 2.821971 × 10^−6^; unfiltered all-sample fit: slope = +4.606595 × 10^−7^, *R*^2^ = 0.002804, *p* = 5.004231 × 10^−2^. Mean shifts remain near zero for all three categories (Fold-switching: +0.000066; IDP: +0.000321; Knotted: +0.000045), indicating only minimal perturbation after region replacement and supporting the main-text Figure 5 interpretation that the blind spot is not dilution-driven.

**Figure S5.**
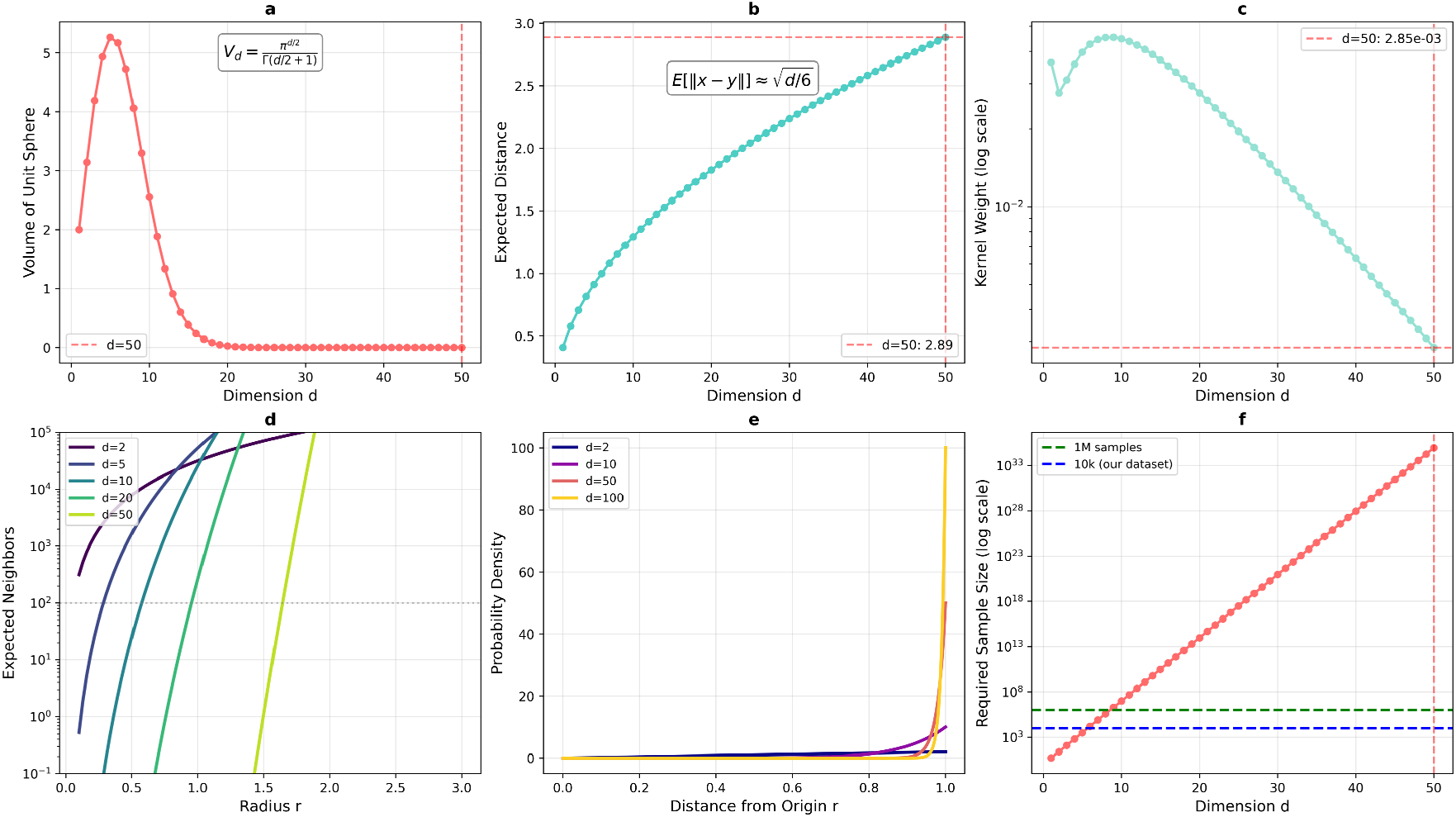
Mathematical foundations of the curse of dimensionality (methodological boundary supporting Figure 6). This figure is not an algorithm benchmark; it derives why local-neighborhood and density notions intrinsically degrade in 50-dimensional spaces from high-dimensional geometry and kernel density estimation (KDE). All panels are based on analytic formulas or controlled numerical evaluation (no empirical fitting). **(a)** Unit-ball volume versus dimension, *V*_*d*_ = *π*^*d/*2^*/*Γ(*d/*2 + 1), rises briefly in low dimensions and then collapses rapidly; at *d* = 50, *V*_50_ = 1.73 × 10^−13^, so a fixed-radius neighborhood covers vanishing effective volume. **(b)** Expected pairwise distance increases with dimension, 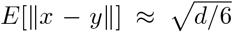 (uniform approximation); at *d* = 50, the expected distance is 2.8868, illustrating distance growth and concentration. **(c)** Gaussian kernel-weight decay under Scott’s bandwidth, *h* = *n*^−1*/*(*d*+4)^*σ* with *n* = 10^4^, and *K*(*x*) = exp(−∥*x*∥^2^*/*2*h*^2^): at *d* = 50, *h* = 0.8432, and the kernel weight at the mean distance is only 2.85 × 10^−3^, so most samples contribute near-zero local density mass. **(d)** Expected-neighbor curves under fixed sample size show severe neighbor scarcity in higher dimensions: for the same radius, effective neighbor counts shrink sharply as *d* increases. **(e)** Concentration-of-measure illustration via radial density *p*(*r*) = *dr*^*d*−1^: probability mass shifts toward the boundary as dimension grows, further weakening near-vs-far discrimination. **(f)** Exponential sample-complexity growth: to keep comparable neighborhood resolution (illustrative base *k* = 5), required samples scale as *k*^*d*^; at *d* = 50, this is 8.88 × 10^34^, i.e., a deficit of 8.88 × 10^30^× relative to *n* = 10^4^. Together, these results imply that in 50D, kernel weights are globally near zero, density estimation becomes quantized/discrete, and correlation analysis becomes unstable.

**Figure S6.**
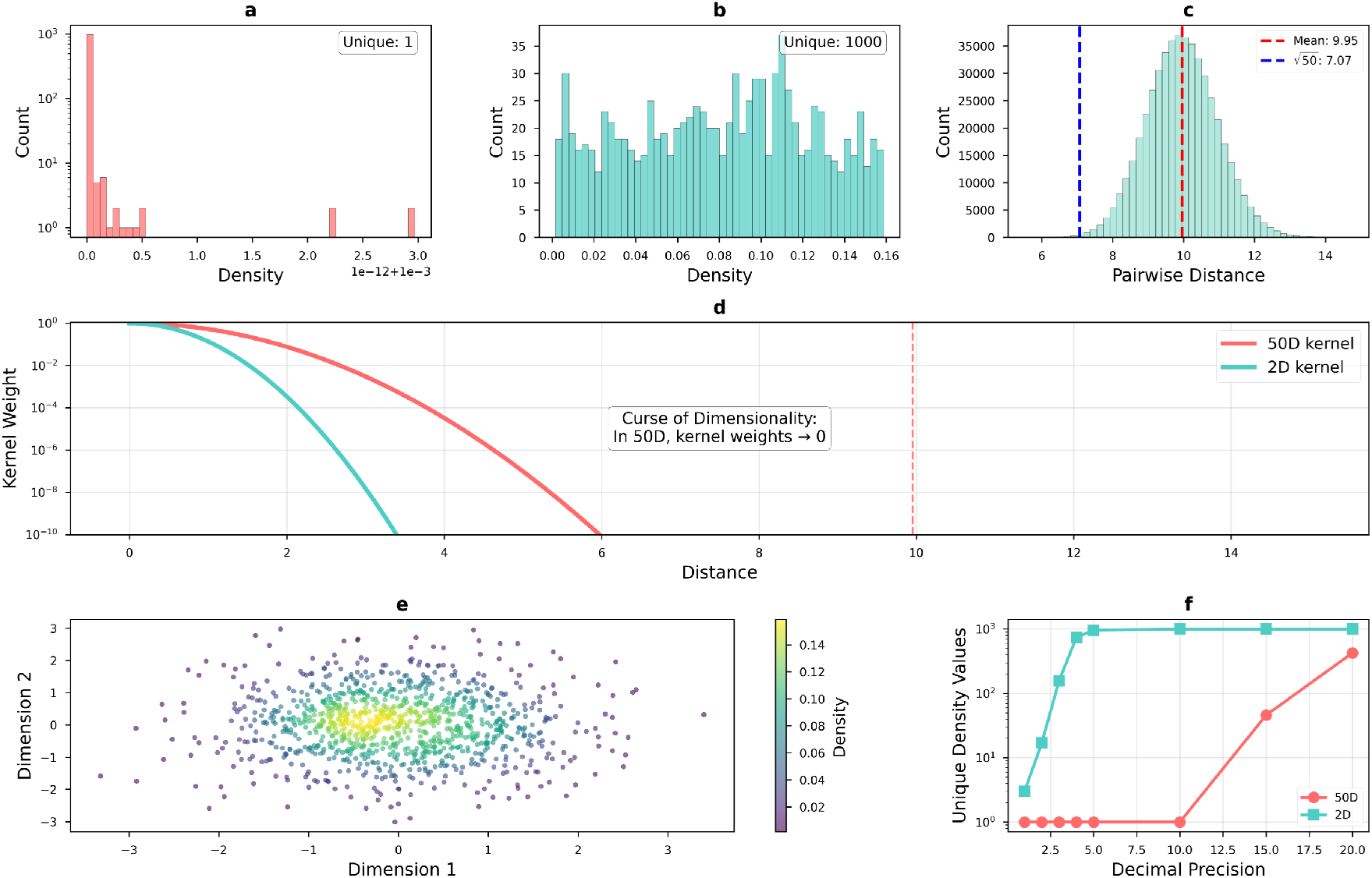
High-dimensional density simulation and quantization collapse. To visualize how kernel density estimation (KDE) degrades in 50-dimensional space, this figure compares density resolution in controlled 50D vs 2D simulations (*n* = 1000 Gaussian samples per setting). **(a)** Histogram of 50D local density values: under Scott’s bandwidth, densities collapse to an almost constant value, yielding only 1 unique value at 10-decimal rounding. **(b)** Histogram of 2D KDE densities: with the same sample size, density remains highly resolvable, yielding 1000 unique values at 10-decimal rounding. **(c)** Pairwise-distance distribution in 50D: mean distance is 9.95 *±* 0.98, close to the theoretical scale 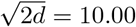, demonstrating strong distance concentration. **(d)** Gaussian kernel-weight decay (50D vs 2D): in 50D, kernel weights decay extremely fast; at the mean distance, the weight is only ∼2.19 × 10^−31^ (Scott bandwidth *σ* = 0.835958), so nearly all local-density contributions are effectively zero. **(e)** 2D scatter with KDE coloring, showing a continuous and resolvable density gradient in low dimension. **(f)** Number of unique density values versus decimal precision: 50D remains low-resolution across precisions, whereas 2D remains high-resolution; at 10 decimals, the resolution ratio is 1000× . Together, these results show that at fixed sample size, high-dimensional KDE undergoes severe quantization collapse, substantially weakening the interpretability of conventional “local density” statistics in 50D.

**Figure S7.**
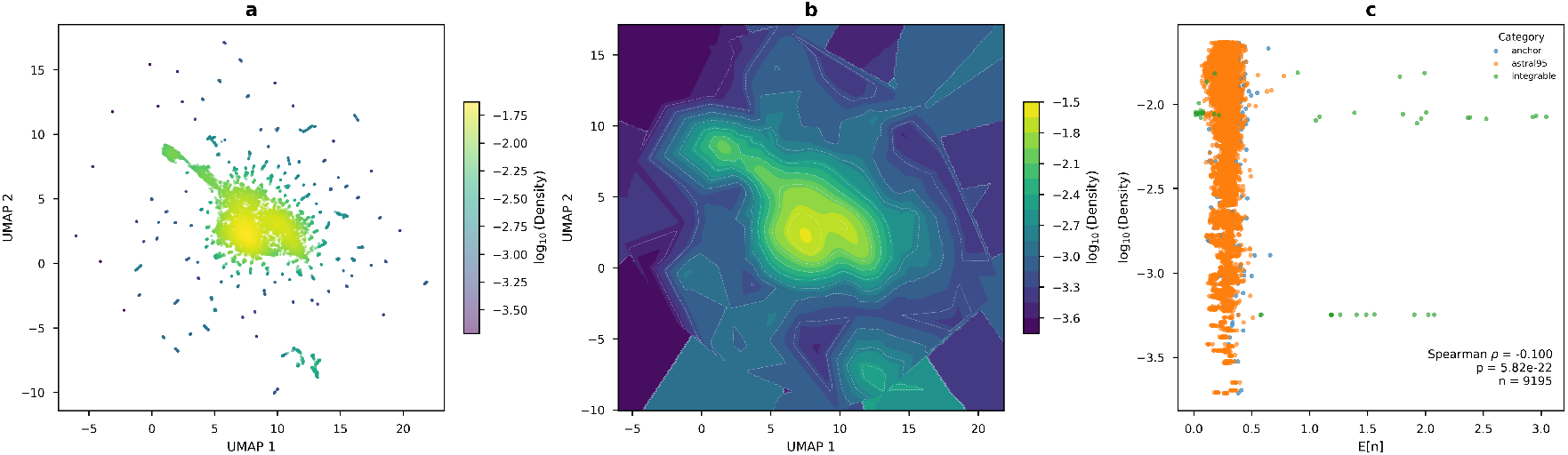
Density landscape and entropy association. Using density estimates and UMAP coordinates for 11,068 protein sequences, this figure visualizes manifold density structure and its statistical association with the entropy-related metric *E*[*n*]. **(a)** UMAP 2D density scatter, where each sample is colored by log_10_(Density), highlighting global density gradients and manifold hotspots. **(b)** Smoothed density contour map over the UMAP plane (contourf + contour): log_10_(Density) is interpolated on a 200 × 200 grid using linear interpolation, with nearest-neighbor filling for boundary holes (NaN fill ratio 34.310%); the value range is approximately [−3.712, −1.631], providing a continuous view of density terrain. **(c)** Scatter plot of *E*[*n*] versus log_10_(Density), colored by category, with *n* = 9,195 valid samples. Spearman correlation is *ρ* = −0.1002 (*p* = 5.818 × 10^−22^), indicating a significant but weak negative association between *E*[*n*] and local manifold density.

**Figure S8.**
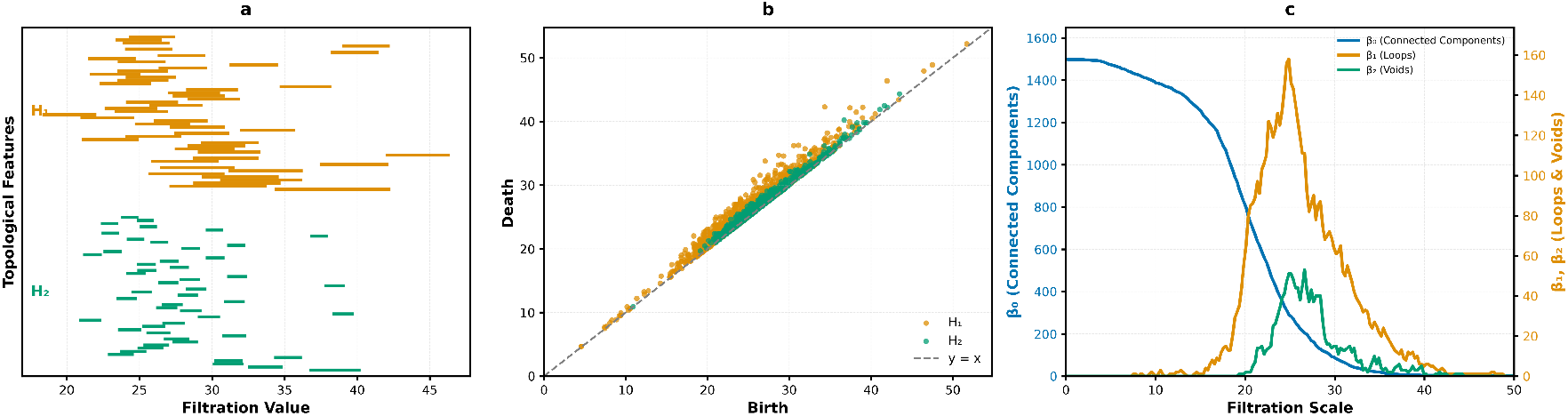
Full-dataset persistent homology overview. Persistent-homology analysis of the full dataset reveals the global organization of mid- and high-order topological structures in the ESM-2 latent space. **(a)** Barcode plot showing only *H*_1_ (loops) and *H*_2_ (voids), ranked by persistence (death−birth), with the top 50 features displayed. This panel highlights the scale span of the most persistent topological features. **(b)** Persistence diagram with *H*_1_/*H*_2_ birth–death scatter and the diagonal *y* = *x*. Valid point counts are *H*_1_ *n* = 1348 and *H*_2_ *n* = 803. Birth/death ranges are [4.5654, 51.6938]*/*[4.6572, 52.1756] for *H*_1_ and [10.8992, 43.4641]*/*[10.9127, 44.3330] for *H*_2_. Persistence summary (median/p90/max) is 0.7716*/*2.4268*/*7.9596 for *H*_1_ and 0.3301*/*1.0042*/*3.5131 for *H*_2_, indicating broader persistence for *H*_1_ than *H*_2_. **(c)** Betti curves (*β*_0_, *β*_1_, *β*_2_) across filtration scale, shown with dual *y*-axes (left: *β*_0_; right: *β*_1_/*β*_2_). Peak values are *β*_0_ = 1500 at scale= 0.0000, *β*_1_ = 158 at scale= 24.8744, and *β*_2_ = 53 at scale= 26.6332. Area under curve over [0, 50] is *β*_0_ = 30459.05, *β*_1_ = 1421.11, *β*_2_ = 348.99. Unlike the bootstrap median*±*IQR summary in main-text Figure 7c, this figure reports the raw topology spectrum from a single full-data computation.

**Figure S9.**
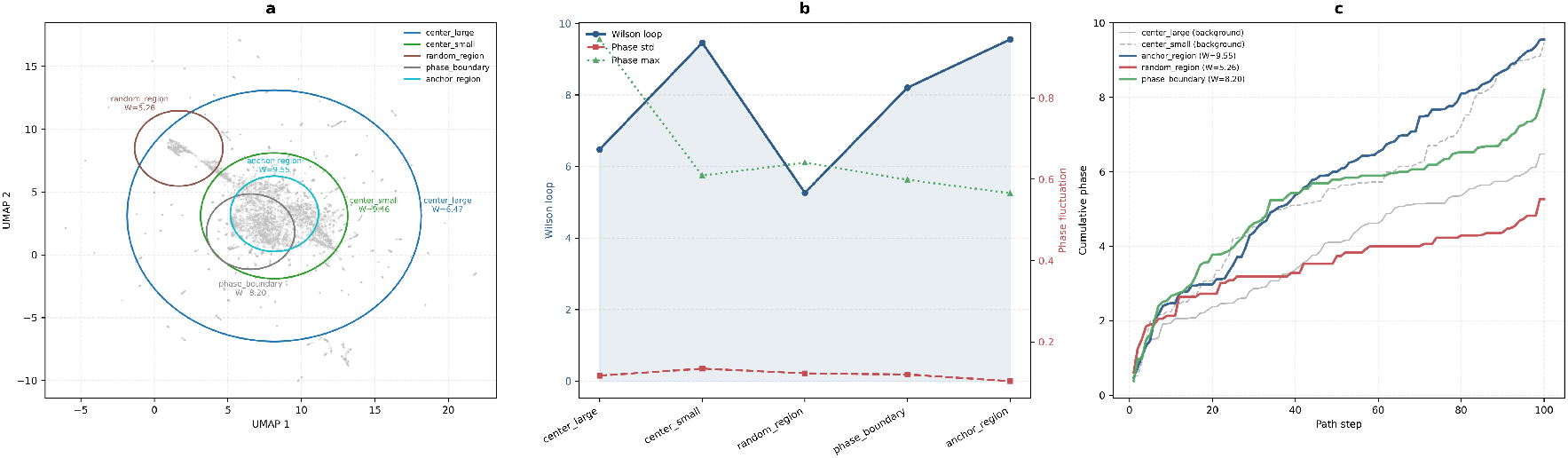
Wilson-loop paths, phase fluctuation, and cumulative phase trajectories. This figure provides a path-level visualization and quantitative comparison of Wilson-loop integrals along five closed loops in the ESM-2 UMAP latent space, complementing main-text Figure 7d (pointwise, microscopic holonomy defect). **(a)** Five explicitly closed paths (center_large, center_small, random_region, phase_boundary, anchor_region) are overlaid on the UMAP 2D background scatter, with per-path Wilson-loop labels. Corresponding Wilson values are 6.4747, 9.4572, 5.2634, 8.2023, and 9.5503. **(b)** A dual-axis line panel compares Wilson loop (left axis) with phase-fluctuation metrics (right axis: std_phase and max_phase). Across paths, Wilson mean/std/range are 7.7896*/*1.6830*/*[5.2634, 9.5503], while phase-fluctuation means are std_phase= 0.1194 and max_phase= 0.6717. Correlations are Corr(*W*, std_phase)= −0.0943 and Corr(*W*, max_phase)= −0.5171, indicating a moderate negative trend between Wilson strength and peak instantaneous fluctuation. **(c)** Cumulative phase trajectories (cumulative phase vs step) highlight three representative paths (highest/lowest/median Wilson: anchor_region, random_region, phase_boundary). Their cumulative endpoints are 9.5503, 5.2634, and 8.2023, respectively, matching their Wilson values one-to-one. The remaining two paths (center_large, center_small) are shown as named background trajectories. This panel emphasizes the integral nature of Wilson loop and visualizes path-dependent differences in accumulation efficiency and fluctuation shape.

**Table S1.** Unified quantitative summary of local compaction and global fidelity/robustness (paired with Figure S3). Local rows report kNN purity (*k* = 15), defined as the fraction of same-class neighbors among each sample’s 15 nearest neighbors, averaged within category; “·/Prevalence” is enrichment = kNN purity / prevalence.

| Scope | Metric / Category | $N$ | Prevalence | PCA | t-SNE | UMAP | PCA/Prev | t-SNE/Prev | UMAP/Prev |
| --- | --- | --- | --- | --- | --- | --- | --- | --- | --- |
| Global | Neighborhood Preservation ($k=15$) | — | — | 0.058 | 0.435 | 0.284 | — | — | — |
| Global | Distance Correlation (Spearman $\rho$) | — | — | 0.909 | 0.674 | 0.777 | — | — | — |
| Global | Seed-mean DC (PCA/UMAP) | — | — | 0.9001 | — | 0.7621 | — | — | — |
| Global | Seed-std DC (PCA/UMAP) | — | — | 0.0079 | — | 0.0150 | — | — | — |
| Global | Variability ratio (UMAP std/PCA std) | — | — | — | — | $1.91 \times$ | — | — | — |
| Global | Mean DC gain (PCA – UMAP) | — | — | +0.1379 | — | — | — | — | — |
| Local | Astral95 | 8292 | 0.7492 | 0.8078 | 0.8570 | 0.8511 | 1.08 | 1.14 | 1.14 |
| Local | Anchor | 856 | 0.0773 | 0.0894 | 0.1180 | 0.1084 | 1.16 | 1.53 | 1.40 |
| Local | Random | 500 | 0.0452 | 0.8604 | 0.9619 | 0.9601 | 19.05 | 21.29 | 21.25 |
| Local | Integrable | 50 | 0.0045 | 0.0533 | 0.2800 | 0.2467 | 11.81 | 61.98 | 54.60 |
| Local | Knotted | 286 | 0.0258 | 0.0692 | 0.1734 | 0.1524 | 2.68 | 6.71 | 5.90 |
| Local | IDP | 1000 | 0.0904 | 0.2703 | 0.7180 | 0.6797 | 2.99 | 7.95 | 7.52 |
| Local | Fold-switching | 84 | 0.0076 | 0.0103 | 0.0254 | 0.0175 | 1.36 | 3.35 | 2.30 |

